# Paired CRISPR/Cas9 guide-RNAs enable high-throughput deletion scanning (ScanDel) of a Mendelian disease locus for functionally critical non-coding elements

**DOI:** 10.1101/092445

**Authors:** Molly Gasperini, Gregory M. Findlay, Aaron McKenna, Jennifer H. Milbank, Choli Lee, Melissa D. Zhang, Darren A. Cusanovich, Jay Shendure

## Abstract

The extent to which distal non-coding mutations contribute to Mendelian disease remains a major unknown in human genetics. Given that a gene’s in vivo function can be appropriately modeled in vitro, CRISPR/Cas9 genome editing enables the large-scale perturbation of distal non-coding regions to identify functional elements in their native context. However, early attempts at such screens have relied on one individual guide RNA (gRNA) per cell, resulting in sparse mutagenesis with minimal redundancy across regions of interest. To address this, we developed a system that uses pairs of gRNAs to program thousands of kilobase-scale deletions that scan across a targeted region in a tiling fashion (“ScanDel”). As a proof-of-concept, we applied ScanDel to program 4,342 overlapping 1- and 2- kilobase (Kb) deletions that tile a 206 Kb region centered on *HPRT1*, the gene underlying Lesch-Nyhan syndrome, with median 27-fold redundancy per base. Programmed deletions were functionally assayed by selecting for loss of *HPRT1* function with 6-thioguanine. *HPRT1* exons served as positive controls, and all were successfully identified as functionally critical by the screen. Remarkably, *HPRT1* function appeared robust to deletion of any intergenic or deeply intronic non-coding region across the 206 Kb locus, indicating that proximal regulatory sequences are sufficient for its expression. A sparser mutagenesis screen of the same 206 Kb with individual gRNAs also failed to identify critical distal regulatory elements. Although our screen did find programmed deletions and individual gRNAs with putative functional consequences that targeted exon-proximal non-coding sequences (e.g. the promoter), long-read sequencing revealed that this signal was driven almost entirely by rare, unexpected deletions that extended into exonic sequence. These targeted validation experiments defined a small region surrounding the transcriptional start site as the only non-coding sequence essential to *HPRT1* function. Overall, our results suggest that distal regulatory elements are not critical for *HPRT1* expression, and underscore the necessity of comprehensive edited-locus genotyping for validating the results of CRISPR screens. The application of ScanDel to additional loci will enable more insight into the extent to which the disruption of distal non-coding elements contributes to Mendelian diseases. In addition, dense, redundant, large-scale deletion scanning with gRNA pairs will facilitate a deeper understanding of endogenous gene regulation in the human genome.

## Introduction

The success of human genetics in identifying the genes and mutations underlying Mendelian diseases has been facilitated by the incontrovertible reality that the majority of causal mutations lie in protein-coding sequences or splice junctions. Indeed, this assumption is explicit in both classic and contemporary practices in genetics (*e.g.* exome sequencing). However, it is clear that distal non-coding mutations make *some* contribution to Mendelian disease. Understanding how often non-coding mutations play a causal role, as well as developing best practices for pinpointing those that do, are critical challenges for the field. For example, in the clinic, even if a patient is diagnosed with a monogenic Mendelian disorder on the basis of phenotype, clinical sequencing mainly of coding regions fails to identify a causal mutation ∼10% of the time (Chong et al., 2015). However, possible explanations include not only distal non-coding mutations, but also misdiagnosis, somatic mutation, technical false negatives, and others.

The picture is very different for the genetics of common disease, where over 90% of disease-associated SNPs fall in non-coding regions (Maurano et al., 2012). Many resources have been developed to predict the location of putative regulatory elements and the effects of regulatory mutations (Ernst & Kellis, 2012; Hoffman et al., 2012; Kircher et al., 2014), with ∼88% of all protein-coding genes tied to a *cis*-expression quantitative locus (eQTL, Aguet et al., 2016), ∼80% of the genome annotated with biochemical function (ENCODE Project Consortium et al., 2012), and numerous tools to link regulatory elements to their target genes (Boyle et al., 2012; Coetzee, Rhie, Berman, Coetzee, & Noushmehr, 2012; Li, Wang, Xia, Sham, & Wang, 2013; Ward & Kellis, 2012). However, the vast majority of these predictions are either confounded (*e.g.* for *cis*-eQTLs, by linkage disequilibrium) or lack functional validation. Indeed, there are few distal non-coding regulatory elements that we can confidently assign to a target gene, or for which we understand the consequences of disruption.

Large-scale functional experiments are clearly an important next step for both common disease genetics (to facilitate the identification of causal regulatory variants and their target genes) and rare disease genetics (to identify distal regulatory elements for Mendelian disease genes where causal non-coding mutations might be found). Within the last year, several studies have used CRISPR/Cas9 genome editing in cell-based screens to introduce and functionally assay large numbers of non-coding mutations at an unprecedented scale (Canver et al., 2015; Chen et al., 2015; Diao et al., 2016; Korkmaz et al., 2016; Rajagopal et al., 2016; Sanjana et al., 2016). The common approach of these studies is to introduce complex libraries of guide RNAs (gRNAs) via lentiviral infection to a population of cells at a low multiplicity of infection (MOI), followed by an assay that queries the function or expression of a gene of interest. CRISPR/Cas9 mediates double-stranded breaks at sites specified by the gRNA in each cell, eventually resulting in a mutation at each targeted site via imperfect non-homologous end joining (NHEJ). A fundamental limitation of these singleton gRNA screens is that because of design constraints (e.g. the uneven distribution of protospacer adjacent motif (PAM) sequences, the variable efficiency of gRNAs, and others), the resulting coverage of regions of interest is incomplete and uneven. As the majority of bases will be perturbed by zero or only one gRNA, these studies rely on the aggregate behavior of clusters of target sites within potential regulatory elements (Canver et al., 2015) or arbitrarily sized windows (e.g., 500 base-pairs) (Sanjana et al., 2016), rather than redundant targeting of each base-pair (bp) by independent gRNAs. Furthermore, it is possible that the mutations introduced by NHEJ at single sites (highly heterogeneous but mainly dominated by small 1-10 bp deletions (McKenna et al., 2016; Tsai et al., 2014)) are insufficient to fully disrupt many regulatory elements.

Here we sought to overcome these weaknesses by introducing *pairs* of gRNAs to each cell, with the goal of inducing a kilobase-scale deletion of the intervening DNA between two programmed cuts. A principal advantage of this method is that by tiling deletions across a region, each targeted base-pair can be covered with high redundancy (<u>scan</u>ning <u>del</u>etion or “ScanDel”). Furthermore, kilobase-scale deletions are much more likely to eliminate the function of an overlapping or fully contained regulatory element, relative to small indels resulting from NHEJ at a single target site. Our approach is analogous to classic deletion scanning experiments (Reid, Gregg, Smithies, & Koller, 1990; Rincon-Limas & Krueger, 1991), but with advantages in throughput and of targeting much larger regions in the endogenous genome rather than sequences cloned to a plasmid.

Adopting the framework of genome-wide CRISPR/Cas9 screens, we synthesized, cloned, and lentivirally delivered thousands of programmed gRNA pairs to cells at a low MOI. Each gRNA pair targets nearby sites, effectively leveraging CRISPR/Cas9’s ability to generate kilobase-scale deletions when NHEJ-mediated repair of two nearby double-stranded breaks results in excision of the intervening DNA segment. In total, we designed and introduced gRNAs pairs programming 4,342 overlapping ∼1- and ∼2- kilobase (Kb) deletions that tiled a 206 Kb region centered on *HPRT1*, the gene underlying the X-linked Mendelian disorder Lesch-Nyhan syndrome. 6-thioguanine (6TG) was used to select for cells that had lost *HPRT1* function. By quantifying gRNA pairs both before and after 6TG selection, we were able to identify programmed deletions that significantly compromised *HPRT1* expression or function.

## Results

### Development of ScanDel

In genome-wide CRISPR/Cas9 screens, a gRNA library is lentivirally delivered to a large pool of cells at a low MOI, such that each infected cell is likely to receive only one gRNA (Shalem et al., 2014; Wang, Wei, Sabatini, & Lander, 2014; Zhou et al., 2014). Each gRNA induces NHEJ-mediated indels centered at the Cas9-mediated cleavage position within the target sequence, with the goal of perturbing the function of the targeted locus. However, given the small and variable length of indels, the robustness of perturbation is inherently limited, particularly when targeting non-coding sequences in which frameshifts are irrelevant. To instead program a kilobase-scale deletion in each cell, we devised the following approach (Fig 1). First, gRNA pairs are designed to program specific deletions (with each gRNA specifying one of the deletion’s boundaries, Fig 1A), and the corresponding pairs of 20 bp spacers are synthesized *in cis* on a microarray (Fig 1B). Second, the paired spacers are inserted into the lentiGuide-Puro plasmid between the U6 promoter and the gRNA backbone. Third, a second gRNA backbone and a second RNA Polymerase (Pol) III promoter (H1 or U6) are inserted between the paired spacers. Fourth, libraries of “gRNA pairs” are lentivirally delivered to a large pool of cells at a low MOI, such that each cell receives a pair of gRNAs that programs a single deletion (Fig 1C). Finally, analogous to conventional genome-wide CRISPR/Cas9 screens, deep sequencing of the integrated gRNA pairs is used as a surrogate measure of the prevalence of each programmed deletion in a population of cells (*e.g.* before and after the cells have been subjected to functional selection) thus capturing the phenotypic consequences of individual deletions.

**Figure 1.**
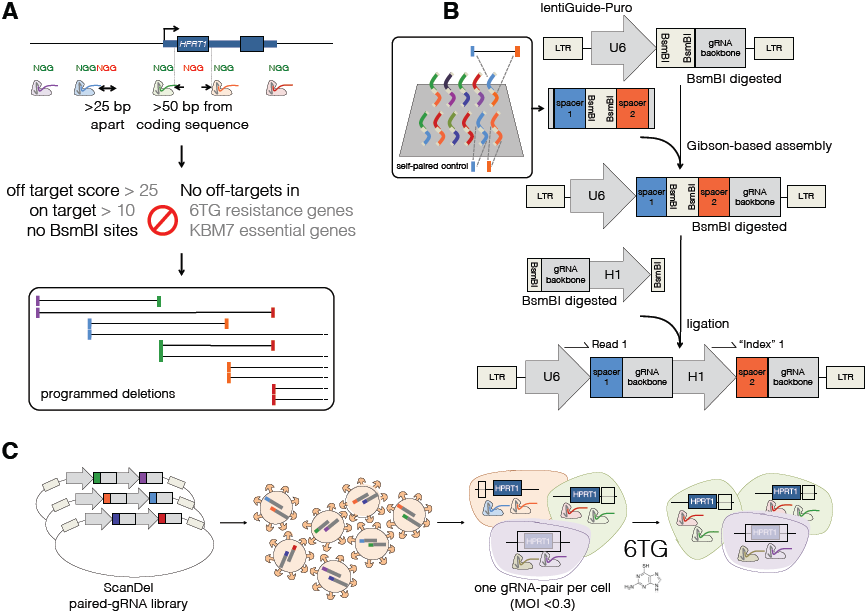
Design, delivery, and selection of ScanDel library of programmed deletions for identification of non-coding regulatory elements. **A)** gRNA pairs are designed from a filtered set of protospacers from all Cas9 PAM sequences (5’-NGGs) in the *HPRT1* locus (see also Fig. 2A). Sites that are >25 bp apart and >50 bp away from exons with on-target efficiency and off-target scores above thresholds are kept. Any spacers with BsmBI restriction enzyme sites or predicted to have off-target hits in other 6TG resistance genes or in KBM7 essential genes (the HAP1 parental cell line) are excluded. Tiles are designed by pairing each remaining spacer to two downstream spacers targeting sequence ∼1 Kb away and ∼2 Kb away. This results in high redundancy of independently programmed, overlapping deletions across the locus (see also Fig. 2B). **B**) All spacer pairs that correspond to programmed deletions are synthesized on a microarray (*inset*). Each spacer is also synthesized as a self-pair as a control for its independent effects. If a self-paired spacer scores positively in the screen, any pairs that use that spacer are removed from analysis (Fig. S2). U6 and gRNA backbone sequence flank the spacer pairs for Gibson-mediated cloning into lentiGuide-Puro (Sanjana, Shalem, & Zhang, 2014), and mirrored BsmBI cut sites separate the spacer pairs to facilitate insertion of a second gRNA backbone and the H1 promoter (*beige*). In the final library, each gRNA is expressed from its own PolIII promoter. This design facilitates PCR and direct sequencing-based quantification of gRNA pair abundances. **C)** The lentiviral library of gRNA pairs is cloned at a minimum of 20x coverage (relative to library complexity) and transduced into HAP1 cells stably expressing Cas9 (via lentiCas9-Blast (Sanjana et al., 2014)) at low MOI. After a week of puromycin selection, the cells are sampled to measure the baseline abundance of each gRNA pair. The final cell population is harvested after a week of 6-thioguanine (6TG) treatment, which selects for cells that have lost HPRT enzymatic function. The phenotypic prevalence of each programmed deletion is quantified by PCR and deep sequencing of the gRNA pairs before and after selection.

As an initial test of our paired guide system, we compared the efficacy of using two different promoters for the two guides (a ‘U6-H1’ system) versus using two copies of the same promoter (‘U6-U6’). We tested these lentiviral gRNA pair expression constructs by targeting the same genomic site for deletion with each system (Supplementary Fig. 1). PCR amplification of the site was performed with unique molecular identifiers (UMIs) in order to minimize biases related to amplicon size (see Methods). The U6-H1 system induced more programmed deletions than the U6-U6 system (20% vs. 10% of reads from cells one week after transduction). The U6-H1 system has several advantages (*e.g.* avoiding recombination between the two U6 promoters during cloning; unique primer design for deep sequencing of each gRNA), and we therefore proceeded with it.

An important caveat for ScanDel, relative to conventional gRNA cell-based screens, is that deletions programmed by gRNA pairs only occur in a minority of cells (Byrne, Ortiz, Mali, Aach, & Church, 2015; Canver et al., 2014), with the other major outcomes being small NHEJ-mediated indels at one or both gRNA-targeted sites. For example, in our test of the U6-H1 system, 32% of edited cells contained the programmed deletion, while the remainder were mutated at one or both gRNA-targeted sites but retained the intervening sequence. While this complicates interpretation, the problem can be overcome by using a robust functional assay in conjunction with multiple, independent gRNA pairs that query the same genomic region, as well as by including unpaired gRNA controls to ensure that observed effects do not occur with individual gRNAs (but rather are dependent on the presence of both gRNAs).

### Application of ScanDel to survey a 206 Kb region surrounding *HPRT1*

With the goal of investigating the potential of non-coding mutations to compromise its function, we applied ScanDel to a 206 Kb region on the X chromosome centered on the *HPRT1* gene (Figs. 1A & 2A). *HPRT1* is a broadly expressed housekeeping gene that encodes the enzyme hypoxanthine(-guanine) phosphoribosyltransferase (HPRT). Loss-of-function mutations in *HPRT1* result in Lesch-Nyhan syndrome (Lesch & Nyhan, 1964), in which a minority of patients present with reduced HPRT enzymatic activity despite the absence of coding mutations (Fu et al., 2014). Such individuals could carry unidentified non-coding mutations that result in reduced *HPRT1* expression. Loss of *HPRT1* also causes resistance to the drug 6TG, a purine analog and chemotherapeutic agent. Thus, it is straightforward to assay cell populations for loss of *HPRT1* function, as only cells with highly reduced expression of functional HPRT will survive selection by 6TG (Fig 1C).

**Figure 2.**
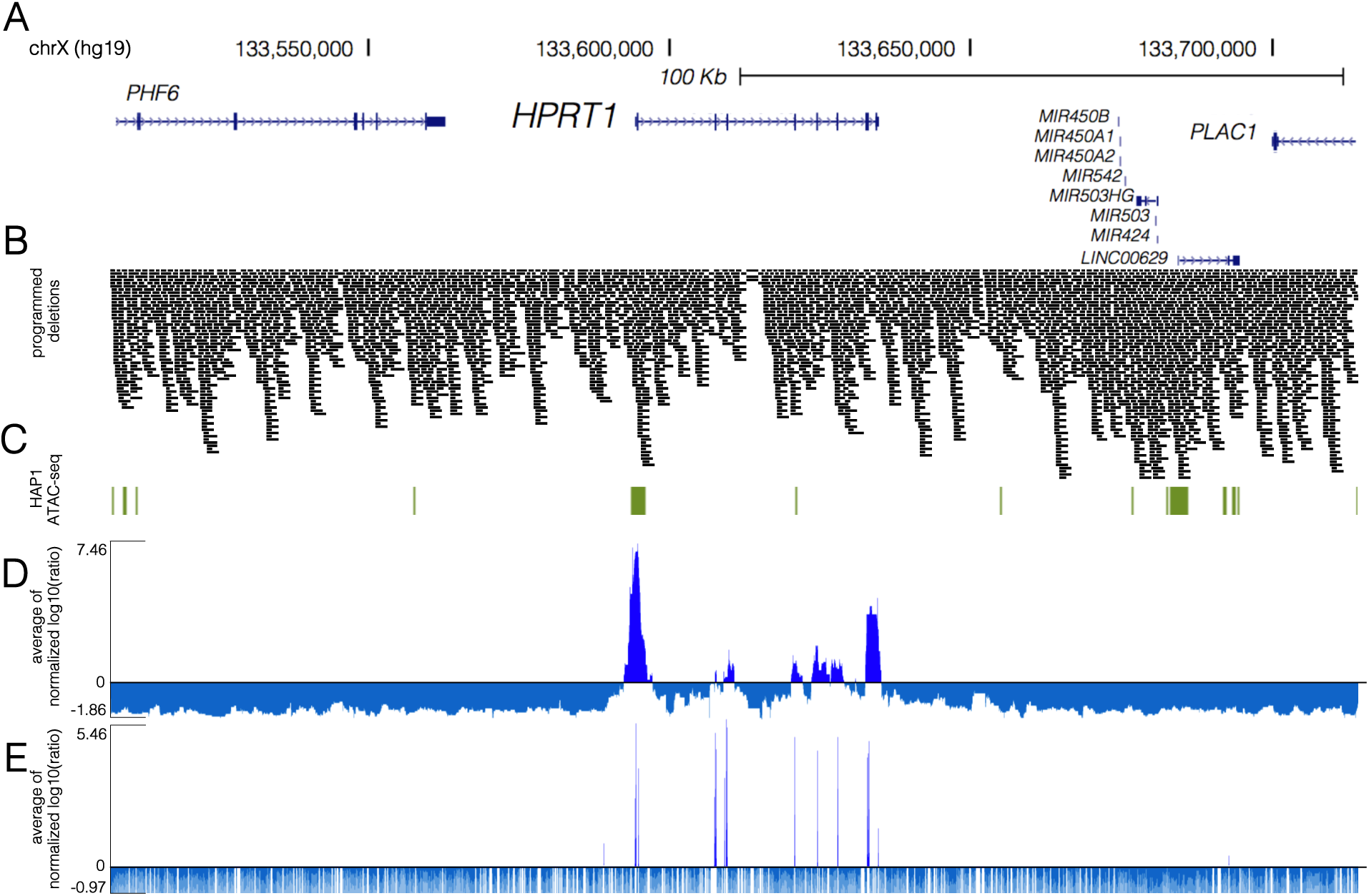
High coverage ScanDel library across the *HPRT1* locus reveals a paucity of critical distal regulatory elements. **A)** Deletions were programmed across 206.1 Kb of the *HPRT1* locus and its surrounding sequence (chrX:133,507,694-133,713,798, hg19, UCSC Genes track in blue). Disruption of *HPRT1* causes Lesch-Nyhan, a neurogenetic Mendelian disorder. **B**) A total of 4,342 1 Kb or 2 Kb deletions were programmed (see Fig. 1A), tiling across the locus such that each base-pair was interrogated by a median of 27 independently programmed deletions. A high density of repeat elements results in reduced coverage of a region within *HPRT1’s* intron 3. Deletions are visualized as black bars spanning the gRNA pair’s programmed cut sites. **C**) HAP1 ATAC-seq hotspots (green) indicate regions of open chromatin in the cell line. Of note, a hotspot extends 600 bp upstream and 1.6 Kb downstream of exon 1. **D**) ScanDel scores were assigned to each base-pair as the average of all selection scores (log10(after/before)) for gRNA pairs that programmed deletions that span that base-pair (Methods). If a gRNA pair used a spacer that was positively selected on its own as a self-pair, the gRNA pair was removed from analysis. Given that depleted gRNAs are usually completely absent after 6TG, their negative scores are of arbitrary negative magnitude. To avoid over-weighting negative values, a minimum score was determined from each replicate’s gRNA pair score distribution (Supplementary Fig. 9), and scores below it were set at this minimum. For each biological replicate, the base-pair’s score was normalized to the replicate’s median of positive scores. The average of the two biological replicates’ normalized scores for that base-pair is displayed, with positive scores in royal blue and negative scores in blue-grey. **E**) An individual gRNA mutagenesis screen of the same region was also performed covering only ∼70% of bases in the region due to the sparsity of high-quality designable spacers. Individual base-pairs were scored based on nearby cut-sites, under the assumption that each gRNA queries a ∼10 bp region. The plotted scores were calculated as in **D**, with positive scores in royal blue and negative scores in blue-grey.

We designed pairs of gRNAs that programmed deletions tiling across the 206 Kb region, including tiles that overlapped *HPRT1* exons in order to allow coding regions to serve as positive controls. As deletion length has been shown to affect deletion rate (Canver et al., 2014), deletions were programmed to be consistently either ∼1 or ∼2 Kb in length (Fig 1A). This design resulted in 4,342-programmed deletions that tiled across the region, collectively covering each base-pair a median of 27 times (Fig 2B). Testing each base-pair with numerous independently programmed, tiling deletions is expected to reduce noise and also increase resolution (as all successfully made deletions tiling a critical regulatory element should exhibit positive selection). However, to guard against the possibility that individual gRNAs’ effects could confound analysis (*e.g.* via off-target mutations, or on-target small ∼10 bp indels), we also included all spacers in the library as pairs with themselves (‘self-pairs’; Fig. 1B **inset**, Supplementary Fig. 2). Additionally, we included 330 negative control gRNA pairs not expected to survive 6TG selection, as they program deletions in non-genic regions far from *HPRT1* or use spacers made of random sequence not present in the reference genome (hg19).

The gRNA pair library was array-synthesized, cloned, and delivered via lentiviral infection to HAP1 cells in replicate (Fig. 1B, C). Cell populations were sampled before and after one week of 6TG selection, with PCR amplification and deep sequencing of gRNA pairs to quantify abundance at each time-point. The functional selection score was calculated as the log10 ratio of normalized read counts after selection relative to before 6TG treatment (“selection score” as log10(after/before 6TG)). Positively scoring self-paired spacers were flagged, and gRNA pairs that used these flagged spacers were excluded from further analysis (11% of pairs in replicate 1 and 3% of pairs in replicate 2). To integrate signal from overlapping programmed deletions, we calculated a “per base-pair” metric as the mean of selection scores of all deletions overlapping a given base (Fig. 2D, Methods). This per base-pair score across the *HPRT1* locus was well-correlated between biological replicates (Pearson: 0.708; Supplementary Fig. 3). Importantly, none of the negative-control gRNA pairs that were sampled in each of the two replicates were positively selected in both experiments (Supplementary Fig. 4).

All *HPRT1* exons exhibited strong functional scores, confirming the sensitivity of ScanDel as applied here to detect sequences essential to *HPRT1* function (Supplementary Fig. 5). However, all of the reproducibly positive non-coding signal across the 206 Kb region was immediately proximal to an *HPRT1* exon. This result suggests that there is no distal regulatory element in the 206 Kb region that is essential to *HPRT1* expression in HAP1 cells.

Near exons, non-coding regions exhibiting positive signal did so even when deletions that also overlapped the exons themselves were excluded from the analysis (Supplementary Fig. 5D). This suggested the presence of essential, proximal regulatory sequences. We noted that the positively scoring regions immediately upstream and downstream of the first exon overlapped with a region of open chromatin identified by performing ATAC-seq in HAP1 cells, supporting the region’s role in gene regulation (Fig. 2C, Supplementary Fig. 5A). This observation motivated us to attempt validation experiments for this region, with the goal of directly confirming which deletions of putative regulatory elements were impairing *HPRT1* function (Fig. 3A, E).

**Figure 3.**
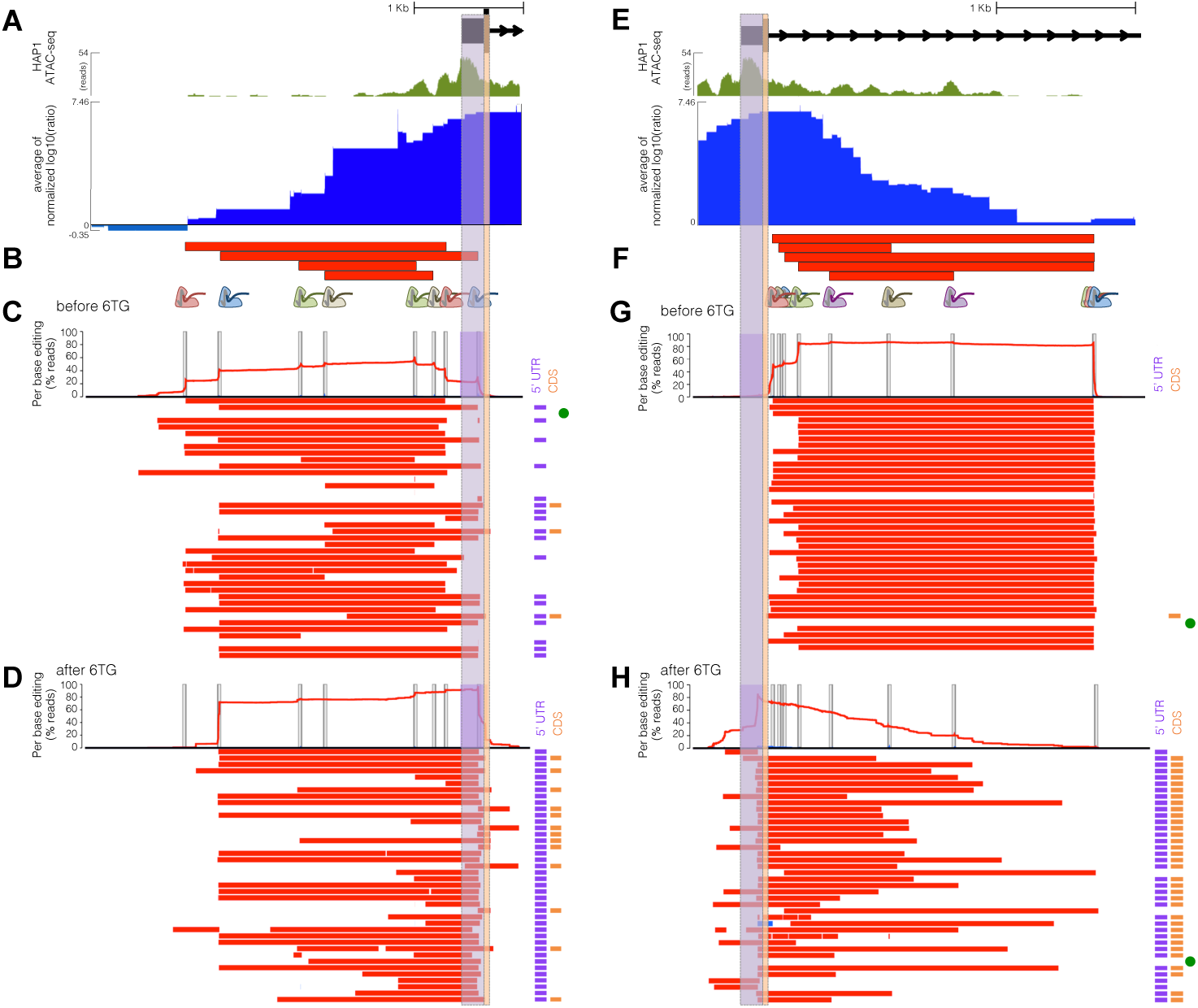
Long-read sequencing of edits derived from exon-proximal ScanDel gRNA pairs reveals rare, unprogrammed, exon-interrupting deletions that drive selective effects. **A)** A putative promoter is implicated by open chromatin (HAP1 ATAC-seq broad peaks, green) surrounding *HPRT1’s* first exon (UCSC Genes, black). ScanDel signal in the 2 Kb upstream of *HPRT1* also suggests the possibility of critical regulatory sequences in this region (blue as in Fig. 2D, chrX:133,591,603-133,594,626, hg19). The 5’ UTR and coding regions of exon 1 are highlighted in purple and orange, respectively. **B)** Four gRNA pairs targeting the promoter were cloned as a small pool, delivered, and selected with 6TG to enable sequencing of the edited locus (programmed deletions displayed as red bars). A 3 Kb region was amplified and sequenced with long reads (Pacific Biosciences). **C)** The chart at the top displays per-base %s for deletions (red) or insertions (blue), with target sites indicated by vertical gray bars. Horizontal bars show the edits found on each haplotype (red: deletions; blue: insertions; ranked by decreasing prevalence). All programmed deletions are abundant before 6TG treatment, in addition to rare, unexpected deletions. The notations to the right indicate if the edits interrupt the TSS/5’UTR (purple bar) and/or coding sequence (orange bar). The unedited haplotype is marked with a green dot. Of note, PCR and sequencing on the PacBio RSII are biased towards smaller fragments, limiting accurate quantitative comparison of read counts from differently sized edits. **D)** Haplotypes from 6TG-selected cells are plotted as in **C**, revealing that only edits that interrupt the TSS/5’UTR survive selection, with no programmed or ‘promoter only’ deletions surviving selection. **E)** Open chromatin (green, as in **A)** and ScanDel signal suggests the presence of critical non-coding regulatory sequences in the first ˜2.7 Kb of intron 1 (chrX:133,593,871-133,596,998, hg19). **F)** 5 gRNA pairs that drove the signal in this intronic region were cloned and 6TG selected as a small pool, as in **C. G)** A 3.1 Kb region spanning the 5’-most part of intron 1 was amplified and sequenced from cells sampled before 6TG selection. Haplotypes and per-base editing rates are diagrammed as in **C. H)** Post-6TG selection haplotypes from the intron 1-targeted cells are plotted as in **G**, revealing that the vast majority of surviving edits disrupt the exon. Two edited haplotypes do not interfere with the exon, but these are present at approximately the level of unedited haplotypes, suggesting 6TG resistance in these cells is caused by mutations elsewhere.

### Direct genotyping of deletions that survive functional selection

With the goal of first validating the positive signal upstream of the first exon, we repeated the experiment with a small pool of 4 gRNA pairs targeting the putative *HPRT1* promoter (Fig 3B). We then amplified 3 Kb of this region by PCR and performed long-read sequencing of the amplicons (Pacific Biosciences). As expected, before 6TG selection, the programmed deletions were all well-represented in the population, although deletions with boundaries deviating from Cas9 cut sites (*i.e.* ‘unprogrammed’) were also detected (Fig 3C). However, after selection on 6TG, deletions with unprogrammed boundaries predominated, including those unseen before 6TG, and that overwhelmingly extend beyond the transcriptional start site (TSS) (Fig 3D). The fact that these initially rare deletions were strongly selected (while 2 Kb promoter deletions that did not cross the TSS were not) suggests that even relatively proximal sequences upstream of the *HPRT1* TSS are not strictly essential for expression. Based on the results of these validation experiments, we conclude that only a narrow window of non-coding sequence upstream of the TSS and 5’UTR is likely relevant to the regulation of *HPRT1* expression.

We next sought to validate the positive signal downstream of the first exon. To do so, we again repeated the experiment with a small pool of just 5 gRNA pairs targeting the first ∼2.7 Kb of intron 1 (Fig 3F). We then amplified the region and again performed long-read sequencing of the amplicons (Pacific Biosciences). As with the promoter, the programmed deletions were all well-represented before 6TG selection, although deletions with unprogrammed boundaries are also detected at a low rate (Fig 3G). After selection, deletions with unprogrammed boundaries predominated again, particularly those that extended into the first exon, thereby disrupting coding sequences (Fig 3H). A low rate of non-exonic deletions survived post-6TG, but these were present at the same level as unedited reads, implying that there may be some other explanation for 6TG resistance in these cells. Thus, as with the promoter, the positive signals that we originally observed for deletions in the first intron were likely consequent to the positive selection of rare ‘on-target-but-with-incorrect-boundaries’ deletions that extend into the first *HPRT1* exon.

### An individual gRNA screen of the same region for comparison to ScanDel

We next compared our ScanDel results against a more conventional screen relying on only individual gRNAs (Canver et al., 2015; Chen et al., 2015; Diao et al., 2016; Korkmaz et al., 2016; Sanjana et al., 2016) (Fig 2E). For this, we cloned a second lentiviral library consisting of 12,151 individual gRNAs targeting the same 206 Kb region and assayed HPRT1 function in HAP1 cells as previously. Under the assumption that each individual gRNA potentially disrupts a ∼10 bp region, this experiment at best interrogates ∼70% of bases within the 206 Kb region due to the sparsity of PAM sites (as compared to our coverage of the entire locus at median ∼27-fold redundancy per base-pair with ScanDel). 85.7% of exon-targeting gRNAs were positively selected (Supplementary Fig. 5) and exonic selection scores were well correlated between biological replicates (Pearson: 0.781). Of 612 negative control gRNAs, none that were sampled in each replicate were positively selected in both experiments (Supplementary Fig. 6). In non-coding sequence, scores were poorly correlated between biological replicates, with a paucity of reproducible, positively selected signal (Pearson: 0.156, Supplementary Fig. 7).

Notably, we did observe a greater proportion of positively scoring gRNAs in the vicinity of exons – *i.e.* whereas only 2% of intergenic gRNAs were positively selected, 7.5% of deep intronic (>2 Kb away from an exon boundary) and 20.5% of proximal intronic (<2 Kb from an exon boundary) gRNAs were positively selected (Fig 4A). Given our earlier observation with ScanDel of rare, ‘on-target-but-with-incorrect-boundaries’ that were confounding when targeting near exon boundaries, we next performed similar validation experiments on individual gRNAs that targeted non-coding sequences nearby exons (Fig 4B). We chose 10 gRNAs in the *HPRT1* promoter region (Fig 4C), and repeated the individual gRNA experiment with a small pool of just these 10 gRNAs, again using long reads (Pacific Biosciences) to sequence the locus before (Fig 4D) and after 6TG selection (Fig 4E). Similar to our results with ScanDel in this region, the only mutations that survived 6TG selection were initially rare deletions whose boundaries extended past the TSS and into the 5’ UTR and/or coding sequence (Fig 4D). This result strongly underscores that caution should be exercised in the interpretation of results from CRISPR-based screens of non-coding regions, whether performed with individual gRNAs or gRNA pairs, and the importance of sequence-based validation of edited regions in the context of such screens.

**Figure 4.**
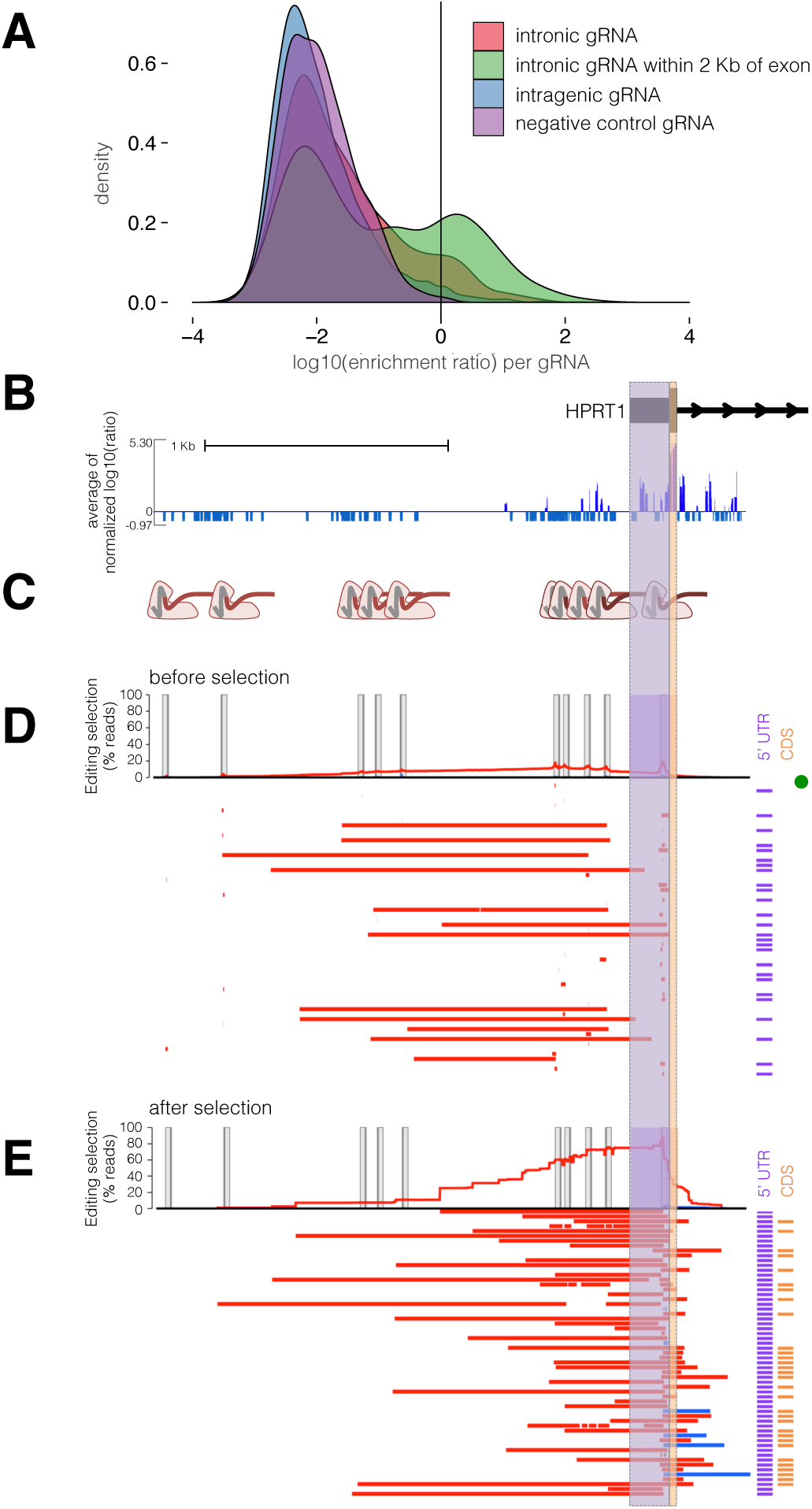
Direct genotyping of edits from an individual gRNA mutagenesis screen also reveals rare, unexpected edits that disrupt *HPRT1*’s exon 1. **A)** A greater proportion of gRNAs targeting non-coding sequence within 2 Kb of exons were positively selected in an individual gRNA screen across the *HPRT1* locus (Fig. 2E, data shown from replicate 1). Each gRNA was assigned a score equal to the log10(after/before 6TG). **B)** gRNAs that target upstream of the transcriptional start site are positively selected. The 2.4 Kb region sequenced for genotype validation (chrX:133,592,240-133,594,646, hg19), *i.e.* a zoom-in of data shown for the whole region in **Fig. 2E. C)** For validation, 10 gRNAs in this 2.4 Kb promoter region were cloned into a low complexity library, delivered to HAP1 cells expressing Cas9, and selected with 6TG. After selection, the 2.4 Kb promoter region was amplified for long-read sequencing. **D)** Before 6TG selection reads are plotted as in **Fig. 3C.** Briefly, the per-base percentage of haplotypes that carried a deletion (red) or insertion (blue) is charted. The edits of the most-prevalent haplotypes from long-read sequencing are drawn as colored bars, and the notations to the right indicate if the edits interrupt the TSS/5’UTR (purple) or coding sequence (orange) of exon 1. A green dot signifies the unedited haplotype. Target site programmed edits are observed and are mainly comprised of the expected small indels, in addition to rarely occurring larger deletions. PCR and sequencing on the PacBio RSII are biased towards smaller fragments, limiting accuracy of quantitative comparison of the read-count prevalence of different sized edits. E) The most abundant haplotypes from cells after 6TG selection are visualized as in D. Only mutations that interrupt exon 1 survive 6TG selection.

## Discussion

We describe a method that uses CRISPR/Cas9 and pairs of gRNAs to experimentally test the functional consequences of thousands of programmed, kilobase-scale genomic deletions in a single experiment. In this proof-of-concept, we introduced a set of densely tiling deletions spanning a 206 Kb region centered on the Mendelian disease gene *HPRT1*, and found no evidence for any distal regulatory element that is critical for its activity, as measured by 6TG sensitivity in HAP1 cells. A screen of this same region with individual gRNAs supported this finding. The dearth of positive selection from disruption of non-coding regions contrasts with the strong positive selection observed from disruption of any exon of *HPRT1*, either by programmed deletions or individual guides. This result is consistent with the relatively small fraction of Lesch-Nyhan patients (0.4%) whose cases go unresolved by clinical sequencing of coding regions (Fu et al., 2014).

*HPRT1* is a widely expressed housekeeping gene (Ardlie et al., 2015) with no eQTLs identified by the Genotype-Tissue Expression Project (Aguet et al., 2016). The simplest explanation of our results is that sequences immediately proximal to the *HPRT1* transcriptional start site may be sufficient to confer the level of expression that provides sensitivity to 6TG, such that even if we disrupt distal regulatory elements that subtly modulate expression, they would go undetected by our strong selection. For future applications of ScanDel, implementing more quantitative readouts will be critical. For example, ScanDel is compatible with any functional selection that reliably separates cells on the basis of gene expression (*e.g.* knocking in GFP to a locus of interest, and then using FACS to stratify ScanDel-edited cells on the basis of expression).

Another possibility, albeit an unlikely one, is that critical regulatory elements for *HPRT1* lie outside of the 206 Kb window that we surveyed. For example, the gene resides at the terminus of a ∼300 Kb topologically associated domain identified in HAP1 cells that spans ∼185 Kb beyond our interrogated region (Sanborn et al., 2015) (Supplementary Fig. 8). This could potentially be addressed by increasing the complexity of the library of programmed deletions in order to densely tile a larger region, or by simply increasing the size of each programmed deletion to interrogate more sequence per gRNA pair.

We note that the paucity of regulatory sequences discovered by CRISPR/Cas9-based screening is not exclusive to this study. Collectively, individual gRNA CRISPR/Cas9 screens have surveyed over a megabase of prioritized non-coding sequences, but only a handful of gRNAs tested have robust phenotypic effects that validate (Canver et al., 2015; Diao et al., 2016; Korkmaz et al., 2016; Rajagopal et al., 2016; Sanjana et al., 2016). One explanation is that the assays being used are insufficiently sensitive and fail to detect modest regulatory effects. This could be addressed through the implementation of more quantitative assays.

A second explanation is that as implemented, genome editing has poor sensitivity due to redundancy in mammalian gene regulation. Redundancy of transcription factor binding sites within enhancers could prevent ∼1-10 bp indels introduced by individual gRNAs from sufficiently disrupting function. Indeed, this was part of the motivation for developing ScanDel, whose programmable kilobase-scale deletions exceed the size of enhancers. Although we did not identify distal enhancers, the essentiality of the TSS and portions of the 5’UTR in our assay was detected primarily by deletions substantially larger than 1-10 bp (Fig. 4D, E), suggesting paired gRNA libraries will be effective for enhancing sensitivity.

A third explanation is that gene expression levels depend in part on historical events, such that disruption of an enhancer in a differentiated cell line would not result in the same outcome as disrupting the same enhancer prior to differentiation. This could be potentially addressed by performing lentivirally-mediated genome editing steps in stem cells, followed by differentiation to a cell type of interest. Any differences in functional consequences that are dependent on the timing of mutation would be of great interest.

Our results also provide a cautionary example of the importance of validation by direct genotyping in the context of CRISPR/Cas9-based screens of non-coding sequences. NHEJ generates a wide assortment of mutations, and strong selections may recover rare editing outcomes. For example, whereas targeting regions adjacent to exons might have been interpreted to reflect the presence of critical proximal regulatory elements, validation experiments using a long-read sequencer showed that this signal was caused by rare deletions that extended into exonic sequence. Many of these unexpected events would have been difficult to detect had we been relying solely on a short-read sequencing platform to genotype editing outcomes. Additionally, validating CRISPR/Cas9-based screens by assessing selection for specific edited haplotypes adds biological information. Here, with long-read genotyping we were able to identify a set of variable deletions that either did or did not drive selection, thus enabling discrimination of essential elements at higher resolution (Fig. 3C, D).

We also note that in experiments relying on pairs (or more) of gRNAs to program deletions, it is critical to include controls that quantify the effects of the individual gRNAs comprising these pairs, as these can have direct effects or off-target effects that might be misinterpreted as being consequent to the programmed deletion. While this manuscript was in preparation, a study was published that similarly used gRNA pairs to program deletion of a large number of lncRNAs, followed by phenotyping for cellular growth (Zhu et al., 2016). Although the results are of great interest, these important controls were not included for the vast majority of spacers used. It will also be important to confirm the validity of each of this screen’s findings through direct genotyping.

Even with the aforementioned open questions and remaining technical hurdles, it is critical that we continue to advance and apply methods for multiplex perturbation of the regulatory landscape with genome editing. The importance of experimental perturbation is highlighted by our results. The non-coding region surrounding *HPRT1’s* first exon resides in open chromatin in this cell line (Fig. 2, Supplementary Fig. 5), yet our results with ScanDel and subsequent validation experiments indicate the essential regulatory region is only a small part of the broader ATAC-seq peak. Perturbing the endogenous genome represents a highly complementary approach to the more classic strategy of reporter assays (Banerji, Rusconi, & Schaffner, 1981; Patwardhan et al., 2009), in which short sequences are tested for their regulatory potential on an episomal vector (of note, the results of early reporter assay-based tests of potential regulatory sequences flanking *HPRT1* are consistent with our findings (Reid et al., 1990; Rincon-Limas & Krueger, 1991)). Indeed, the ongoing challenge that genome editing can address is to understand how short sequences with regulatory potential coordinate with one another across endogenous loci to give rise to specific levels of expression.

In summary, ScanDel enables the multiplex characterization of the functional consequences of thousands of programmed, kilobase-scale deletions to the endogenous genome in a single experiment. We applied ScanDel to *HPRT1*, a housekeeping gene in which disruptive mutations cause Lesch-Nyhan syndrome, introducing densely tiled 1-2 Kb deletions across a 206 Kb region encompassing the gene, covering each base-pair with median ∼27-fold redundancy. Our results demonstrate the absence of distal *cis*-regulatory elements in this region that are critical for *HPRT1* expression. In the future, we anticipate that large-scale perturbation of putative regulatory elements in their endogenous context with methods such as ScanDel will provide further insights into gene regulation and the contribution of non-coding mutations to human disease.

## Methods

### Tissue culture

HAP1 cells were purchased from Horizon Discovery and cultured in Iscove’s Modified Dulbecco’s Medium with L-glutamine and 25 mM HEPES (Gibco). HEK293T cells were purchased from ATCC and cultured in Dulbecco’s Modified Eagle’s Medium with high glucose and sodium pyruvate (LifeTechnologies). Both media were supplemented with 10% Fetal Bovine Serum (Rocky Mountain Biologicals) and 1% Penicillin-Streptomycin (Gibco), and grown with 5% CO_2_ at 37° C.

### gRNA library design

To generate a list of gRNAs, we identified all 20 bp protospacers followed by a 5’-NGG PAM sequence from chrX:133,507,694-133,713,798 (hg19). We then excluded protospacers that had a perfect sequence match elsewhere in the genome, and scored the remaining gRNAs for both on-target and off-target activity. We considered off-target sequences that had five or fewer mismatches to the putative gRNA, and calculated an aggregate off-target score using the method of Hsu et al., 2013 (Hsu et al., 2013). In addition we scored each site for on-target efficiency (Doench et al., 2016). Final deletion pairs were matched using spacers that did not contain BsmBI restriction sites and passed filters as described in Fig. 1. Contrastingly, the individual gRNA library included all of the spacers targeting the same region, excluding those predicted to have 2,000 or more off-targets or to have off-targets with 4 or fewer mismatches within the targeted *HPRT1* region.

### Building the gRNA pair library

This library cloning method was developed in parallel to similar recently published methods (Aparicio-Prat et al., 2015) and is modified from the GeCKO single gRNA cloning scheme (Sanjana et al., 2014; Shalem et al., 2014). First, the lentiGuide-Puro backbone (Addgene #52963) is digested with BsmBI (FastDigest Esp3I, Thermo) and gel purified. The paired spacers (flanked with lentiGuide-Puro overlap sequences) are synthesized twice on a microarray (CustomArray, Inc.) such that each pairing is represented in both possible orders (Supplementary Fig. 2).

To ensure quality of array synthesis, 1 ng of the oligo pool was amplified with Kapa HiFi Hotstart ReadyMix (KHF, Kapa Biosystems) and run on a gel to confirm oligos are of the expected 108 bp length. After PCR purification with Agencourt AMPure XP beads (Beckman Coulter), the amplicon is cloned into lentiGuide-Puro using In-Fusion HD Cloning Plus (Clontech) and transformed into Stable Competent *E*. *coli* (NEB C3040H) to minimize repeat-based recombination of the lentivirus. This ensuing library (lentiGuide-Puro-2xSpacers) now contains each pair of spacers, but is still missing the additional sgRNA backbone and PolIII promoter.

We next cloned in the additional gRNA backbone and H1 promoter between each spacer pairing to enable expression of the two independent gRNAs. The gRNA backbone-H1 promoter fragment was ordered as a gBlock (IDT) with flanking BsmBI sites to allow ligation into the BsmBI-digested lentiGuide-Puro-2xSpacers library. The gBlock and the lentiGuide-Puro-2xSpacers are each digested with BsmBI, purified, ligated together with Quick Ligase (NEB M2200S), and transformed into Stable Competent *E. coli* to create a final lentiGuide-Puro-2xgRNA library.

To prevent bottlenecking of the library, these cloning steps are performed with enough replicates at high efficiency to maintain a minimum of 20x average library coverage (relative to the expected library complexity). Sequencing of the lentiGuide-Puro-2xgRNA library revealed 97.8% retention of diversity from the designed paired spacers. However, 16% of library reads held unprogrammed, interswapped pairs. 88.5% of these swaps are only seen in a single read, implying a more likely cause is template switching during either PCR or cluster generation. For all experimental analysis, only reads of gRNA pairs that perfect matched programmed pairs were considered.

### Building the individual gRNA library

The spacers of this library were similarly synthesized on an array, amplified, and purified as above. The lentiGuide-Puro backbone was linearized as above, and the library cloned into it using the NEBuilder HiFi DNA Assembly Master Mix (NEB). This plasmid was transformed into Stable Competent *E. coli,* generating enough transformants for 30x average coverage. This method produced 98.5% retention of complexity from the designed array.

### Lentiviral library production, delivery, and 6-thioguanine selection

Lentivirus was produced using Lipofectamine 3000 (Life Technologies) to transfect HEK293T with the lentiviral vector libraries made above and 3rd generation packaging plasmids (pMDLg/pRRE Addgene 12251, pRSV-Rev Addgene 12253, pMD2.G Addgene 12259). Supernatant was collected 72 hours after transfection, centrifuged at 300 rcf for 5 minutes to remove cell debris, and passed through a 0.45 μm syringe filter.

To create a monoclonal HAP1 cell line stably expressing Cas9, HAP1 cells were transduced with lentivirus produced using lentiCas9-Blast (Addgene 52962), selected with 5 μg/mL Blasticidin (Thermo Fisher Scientific), and single-cell sorted via FACS.

HAP1-Cas9-Blast monoclonal cells were plated to be at 30% confluency on the day of lentiviral gRNA/pair transduction. To transduce, 5% of the recipient cells’ media was replaced with filtered virus, limiting the multiplicity of infection to < 0.3. Media was changed after 24 hours, and selection for transduced cells began 48 hours post-transduction. Puromycin was added at 2 μg/mL for two days to assess the percentage of cells transduced, and then cells were maintained in 1 μg/mL for 5 more days.

After puromycin treatment, an initial population of cells was collected. Selection for loss of HPRT function was performed by applying 5 uM 6TG to the remaining cells at <50% confluency for 7 days. Enough cells were transduced and sampled at each timepoint to maintain minimum 2,000x average coverage of the library in each population.

Sequencing of the baseline (*i.e.* pre-6TG) population revealed 98.4% of diversity of the lentiGuide-Puro-2xgRNA library was preserved from replicate 1, and replicate 2 retained 78.8%. As our deletions are highly overlapping, we proceeded with replicate 2 as all base-pairs are interrogated despite the lower diversity. We observed 95.6% retention of programmed library diversity in replicate 1 of single gRNA plasmid library and 71.2% of replicate 2.

Interswapped gRNA pairs were observed in 35.5% of reads from the baseline pre-6TG sample. This is an increase from the 16% observed in reads from the lentiGuide-Puro-2xgRNA plasmid library. This suggests additional template switching during the library’s amplification from gDNA, which requires more cycles of PCR. However, since we are directly sequencing each gRNA spacer as a read out opposed to using barcoded libraries (Zhu et al., 2016) and only taking exact sequence matches, this does not pose a problem.

### gRNA library amplification and sequencing from HAP1 cells

gDNA was extracted from the cells sampled before and after 6TG selection using the DNeasy Blood & Tissue kit (QIAGEN). KHF was used for all amplification steps. The libraries were initially amplified from a minimum of 6 ug of gDNA across thirty 50 μL reactions, ensuring sampling of ∼2 million haploid genome equivalents at each timepoint. Two additional PCRs were performed to add sequencing adapters and sample indices to the amplicon, with AMPure bead purification between each reaction. Amplification conditions were optimized using qPCR to minimize overamplification of the construct.

Sequencing was performed on an Illumina Miseq using a 50-cycle kit. Read 1 and the Illumina Index read were used to sequence the two gRNAs in the paired gRNA construct prior to paired-end turnaround, and Read 2 was used to sequence the 9 bp sample index.

### Calculation of a selection score assignment per base-pair

Custom Python scripts counted tallies of gRNAs (for individual gRNA library experiments) or gRNA pairs before and after selection. These counts were normalized to the total number of reads per sample. An enrichment ratio was calculated for each gRNA/pair by dividing its normalized read count after selection by its before selection read count. A selection score is the log10 of the enrichment ratio (log10(after/before)). If a gRNA or gRNA pair was absent before selection, it was excluded from further analysis. Any gRNA pairs that used a self-paired gRNA with an independent selection ratio > 0 were also excluded from further analysis.

If a gRNA/pair is absent after 6TG selection, its selection score as calculated will be a negative number relatively large in magnitude that is somewhat arbitrarily determined by the number of pre-selection reads. Thus, to limit the contribution of these scores to average measurements derived from many independent deletions, we set a minimum selection score equal to the middle of the bimodal distribution between the positively and negatively selected deletions of each replicate (Supplementary Fig. 9). For example, in ScanDel replicate 1, if the log10-value of a selection score was less than −0.35, that gRNA pair’s score was set to −0.35. Each individual base-pair was assigned a per base-pair selection score by taking the mean of all deletions programmed to cover that base-pair. The per base-pair score was normalized to the median score for all positive scores in that replicate. The per base-pair selection score of each replicate was averaged to get the final selection score per base-pair. Per base-pair scores were uploaded as a bedgraph for visualization on the UCSC Genome Browser (http://genome.ucsc.edu).

For the individual gRNA mutagenesis screen, we calculated selection scores per base-pair similarly, assuming a 10 bp deletion was made by each gRNA queried. If a base-pair was scored at the minimum negative threshold in one screen, it was given that value for the consensus selection score of the two replicates.

### Bulk ATAC-seq of HAP1 cells

Two biological replicates were separately maintained (on 10cm dishes, split 1:10 three times per week) and processed separately. Chromatin accessibility in the HAP1 cell line was profiled with the ATAC-seq protocol (Buenrostro, Giresi, Zaba, Chang, & Greenleaf, 2013) with slight modifications. The media for 10cm plates of confluent HAP1 cells was aspirated and replaced with 2 mL of ice cold lysis buffer (‘CLB+’; made as described in the original paper, but supplemented with protease inhibitors (Sigma cat. no. P8340)). Cells were incubated on ice for 10 minutes in CLB+ and then were dislodged with a cell scraper and transferred to a 15 mL conical tube and pelleted at 500 rcf for 5 min at 4° C. Nuclei were resuspended in 1 mL of CLB+ and counted on a hemocytometer. 50,000 nuclei in 22.5ul of CLB+ were combined with 2.5 μL of TDE1 enzyme and 25ul of TD buffer (Illumina). Tagmentation conditions were as described in the original paper (37°C for 30 min). After MinElute purification into 10 μL EB buffer (Qiagen), 5 μL of tagmented DNA was amplified in 25 μL reactions for 12 cycles using the NEBNext Master Mix (NEB). Reactions were monitored with SYBR Green to ensure that samples were not overamplified. PCR products were cleaned once with a QiaQuick PCR Cleanup Kit (Qiagen) and once with 1x AMPure beads (Agencourt). The quality of the library was assessed on a 6% TBE gel and the yield was measured by Qubit (1.0) fluorometer (Invitrogen).

Samples were sequenced on two paired-end Illumina NextSeq 500 runs. Read lengths were 2x75 bp for the first run and 2x151bp for the second run, so the second run was truncated to 75 bp. Sequencing reads were also trimmed for read-through of adapter sequences and quality with Trimmomatic ((Bolger, Lohse, & Usadel, 2014),‘NexteraPE-PE.fa:2:30:10:1:true TRAILING:3 SLIDINGWINDOW:4:10 MINLEN:20’ parameters) and then mapped to the 1000 genomes integrated reference genome ‘hs37d5’ (ftp://ftp.1000genomes.ebi.ac.uk/vol1/ftp/technical/reference/phase2_reference_assembly_sequence/>) with bowtie2 (Langmead & Salzberg, 2012), using the ‘-X 2000 −3 1’ parameters. Only properly paired and uniquely mapped reads with a mapping quality above 10 were retained (‘samtools -f3 -F12 -q10’). Reads mapping to the mitochondrial genome and non-chromosomal contigs were also filtered out. In addition, duplicate reads were removed with Picard (http://broadinstitute.github.io/picard/). After checking QC metrics on the individual replicates, reads from the two libraries were combined for downstream analysis. Hypersensitive sites were called (at a 1% false discovery rate) with the Hotspot algorithm (John et al., 2011).

### Validation and direct genotyping of positive signal from the screens

gRNA pairs that drove the ScanDel signal surrounding *HPRT1’s* first exon were cloned into simple lentiGuide-Puro-2xgRNA libraries. The TSS ScanDel validation library contained four pairs and the intron 1 library contained five (Supplementary Tables 1–2). For the individual gRNA screen TSS library, ten gRNAs were cloned into lentiGuide-Puro (Supplementary Table 3). These constructs were lentivirally delivered to HAP1-Cas9-Blast cells, selected with 6TG, and gDNA extracted as described above.

As the expected deletions could remove up to two kilobases, the loci were sequenced with a Pacific Biosciences RSII (University of Washington PacBio Sequencing Services, P6C4 chemistry, RSII platform). To prepare libraries for PacBio sequencing, the TSS- or intron 1-targeted regions were amplified from 800 ng of gDNA each, using four 50 μL KHF reactions with primers adding sample indices and SbfI or NotI cut sites. The purified amplicons (Zymo Research DNA Clean & Concentrator-5) were digested with SbfI-HF (NEB) and NotI-HF (NEB), leaving sticky ends. 5’-phosphorylated SMRT-bell hairpin oligos (IDT) containing the PacBio priming site, hairpin-forming sequence, and resulting sticky ends for either SbfI or NotI were annealed by heating to 85° C and snap frozen in 10mM Tris 8.5, 0.1mM EDTA, 100 mM NaCl. These were ligated at 10x molar excess to the digested amplicons, destroying the restriction site once attached. To remove undigested amplicons and primers, this ligation was performed in the presence of further SbfI and NotI, and followed by treatment with Exo7 (Affymetrix) and Exo3 (Enzymatics).

Only reads with over five circular consensus sequence passes and containing the expected first twelve 5’ and 3’ base-pairs of the amplicon were used for further analysis. Reads positive for complex inversions (>= 100 basepairs) were removed from the library using the Waterman-Eggert algorithm with match, mismatch, gap open, and gap extend scores of 2, 10, 10, and 5, respectively (Döring, Weese, Rausch, & Reinert, 2008). The resulting reads were then were aligned to the amplicon reference using the NEEDLEALL (Rice, Longden, & Bleasby, 2000) aligner with a gap open penalty of 10 and a gap extension penalty of 0.5. Insertions were required to start within a window of five bases up or downstream of the putative cut site. Deletions were required to either start or end within the same 10 bp window or span the window. Reads that carried the same edit pattern were collapsed into haplotypes, and figures were generated using a custom D3 script.

### Comparing deletion rate of U6-H1 versus U6-U6

Two protospacers were chosen to program a 365 bp deletion within the second intron of *HPRT1* and their spacers were cloned into a U6-H1 construct and U6-U6 construct (Supplementary Fig. 1). Virus was produced and delivered to cells, which were selected with puromycin, and gDNA extracted as described above. The locus was amplified in four successive rounds of nested PCR. The first reaction was only 3 cycles and included a forward primer with a 10-bp unique molecular index (UMI). The second reaction amplified any UMI-tagged fragments. The third and fourth reactions added sample indices and Illumina flow cell adapters. The products were AMPure cleaned between each reaction at a concentration that would lose primer dimer but retain the smaller deletion-holding fragments, and sequenced on a MiSeq. Any reads that contained the same UMI or edit pattern were collapsed using custom scripts and their alignments were visualized with the same D3 script as above.

## Acknowledgements

For discussion and advice, the authors thank all the members of the Shendure Lab, particularly Lea Starita, Andrew Hill, Ron Hause, Seungsoo Kim, Martin Kircher, and Beth Martin. We thank the University of Washington PacBio Sequencing Services core for their assistance. This work was supported by an NIH Director’s Pioneer Award (JS; DP1HG007811), National Human Genome Research Institute (NHGRI) (JS; 1R01HG006768), NHGRI and Division of Cancer Prevention, National Cancer Institute (JS; 1R01CA197139). MG is a National Science Foundation Graduate Research Fellow. DAC was supported in part by T32HL007828 from the National Heart, Lung, and Blood Institute. JS is an investigator of the Howard Hughes Medical Institute.

## Author contributions

MG, GMF, and JS developed the initial concept. MG led experiments and analysis, together with GMF and with assistance from JHM and MDZ. AM performed guide on/off target scoring and development of Cas9 edit visualization pipeline. DAC performed and analyzed ATAC-seq in HAP1 cells. CL designed and implemented the custom sequencing protocol. MG, GMF, and JS designed experiments, interpreted data, and wrote the manuscript. JS provided overall supervision.

---

**Supplementary Figure 1:**
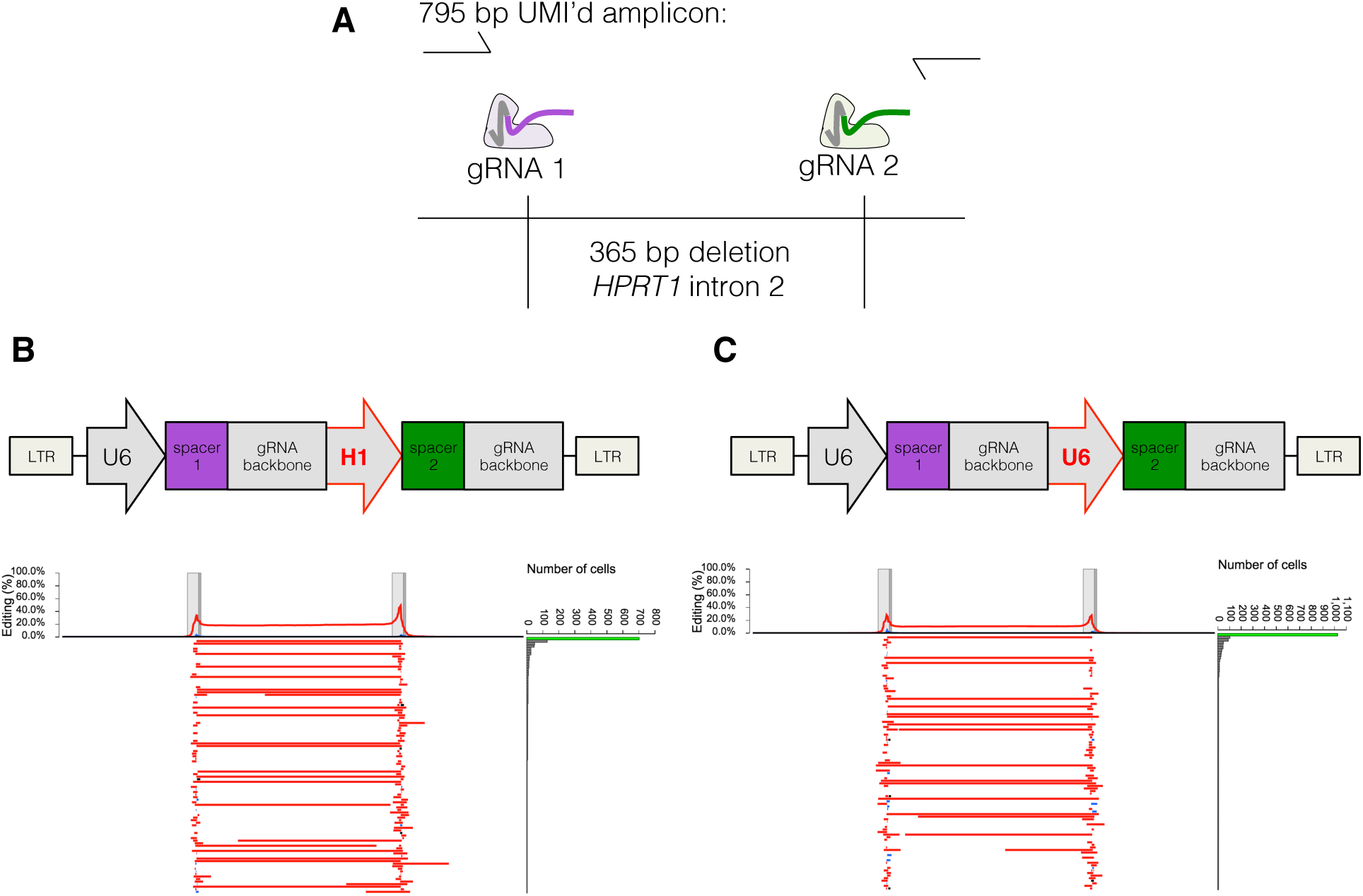
The U6-H1 gRNA pair expression construct induces a higher deletion rate. **A)** Two spacers were chosen to program a 365 bp deletion within the second intron of *HPRT1.* To test deletion efficiency of the method as described in Fig. 1C, virus was made from the constructs depicted in **B** and **C**, and separately transduced into HAP1 at MOI < 0.3. Following 1 week of puromycin selection, gDNA was extracted and the targeted region amplified. The first 3 cycles of this PCR contained a forward primer with a unique molecular tag (UMI) to track reads from the same original cell. Sequencing was performed on a MiSeq. Of note, PCR bias for smaller deletion-holding amplicons was reduced by collapsing reads with the same UMI, but the potential remains for higher clustering efficiency of the shorter amplicons. **B)** The spacers for the deletion in **A** were placed behind either a U6 or H1 PolIII promoter. 20% of sampled haplotypes contained the programmed deletion, but 36% of sampled haplotypes remained unedited, implying longer editing time could result in a higher deletion rate. Reads were generated as described in **A**, and aligned as described in Methods and Fig. 3. The per base-pair editing rate summed across all sampled haplotypes is charted as a percentage at top, and the top 100 most prevalent haplotypes are displayed below it. Red indicates deletions and blue insertions. **C)** The spacers for the deletion in **A** were each placed behind a U6 PolIII promoter, and delivered, sampled, and visualized as above. With this expression construct, 10% of sampled haplotypes contained the programmed deletion.

**Supplementary Figure 2:**
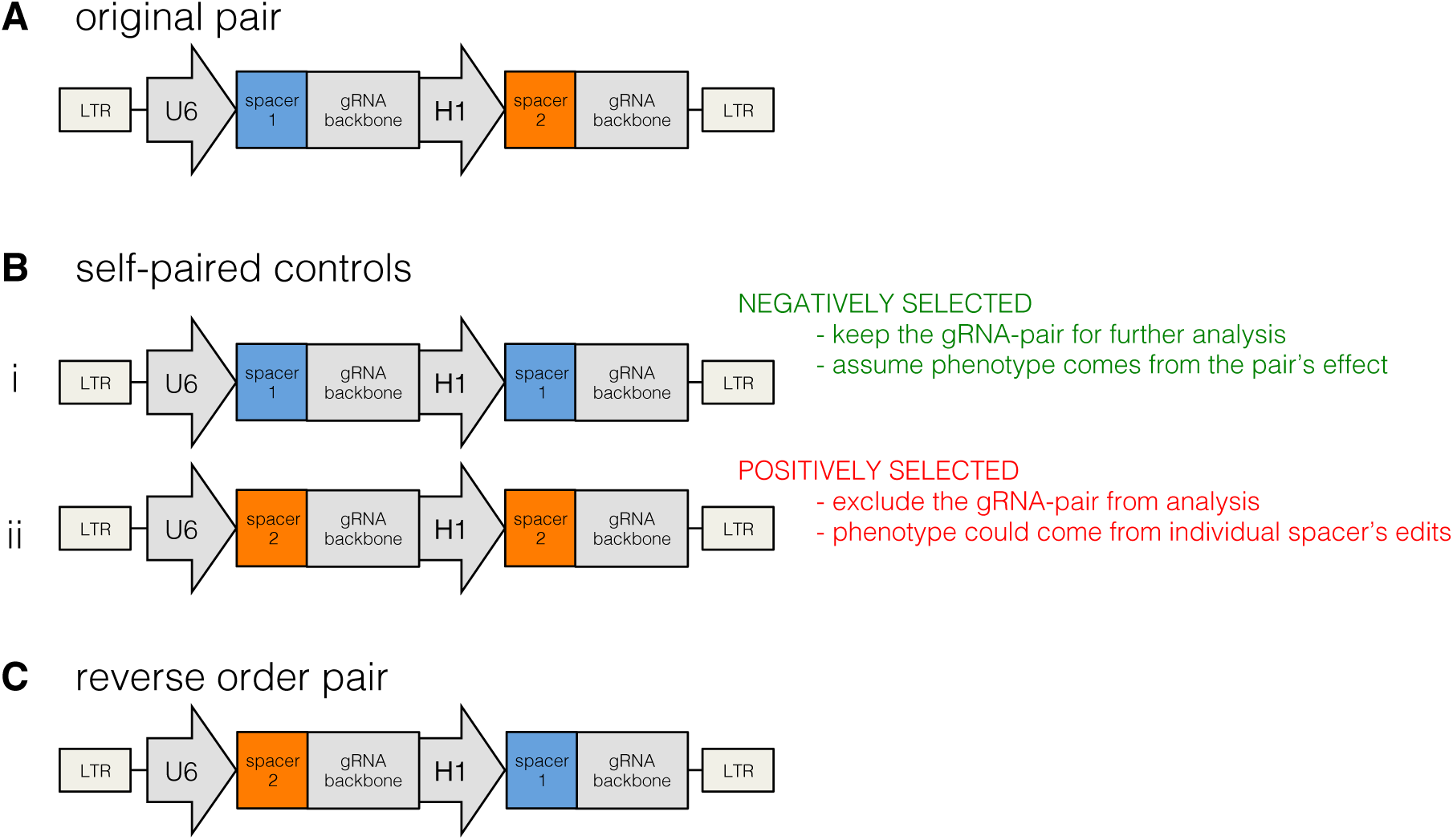
Self-paired spacers in the ScanDel library reveal phenotypes independently created by individual spacers. **A)** The spacers used in every designed gRNA pair had their own self-paired control included in the programmed gRNA pair library. **B)** The self-paired controls consisted of the exact same spacer included behind each promoter in the expression construct (two for each pair; (*i*) and (*ii*)). If a self-paired spacer was positively selected, any gRNA pairs that included that spacer were excluded from further analysis. This avoided any confounding effects of alternative repair outcomes that result from an individual gRNA’s edit that could cause 6TG resistance (*e.g.* a ∼10 bp indel disrupting a transcription factor binding site, or disrupting an off-target locus that affects 6TG resistance, or an individual gRNA inducing translocations of *HPRT1* at a high rate). By excluding these gRNAs, we can more confidently attribute observed phenotypes to programmed deletion induced by the gRNA pairs. **C)** Each gRNA pair was included in both possible orderings on the microarray. This was intended to minimize the impact of differences between the promoters, as well as to increase the chance that each deletion will be represented in the library, as synthesizing each pair twice reduces loss due to synthesis errors and cloning bottlenecks.

**Supplementary Figure 3:**
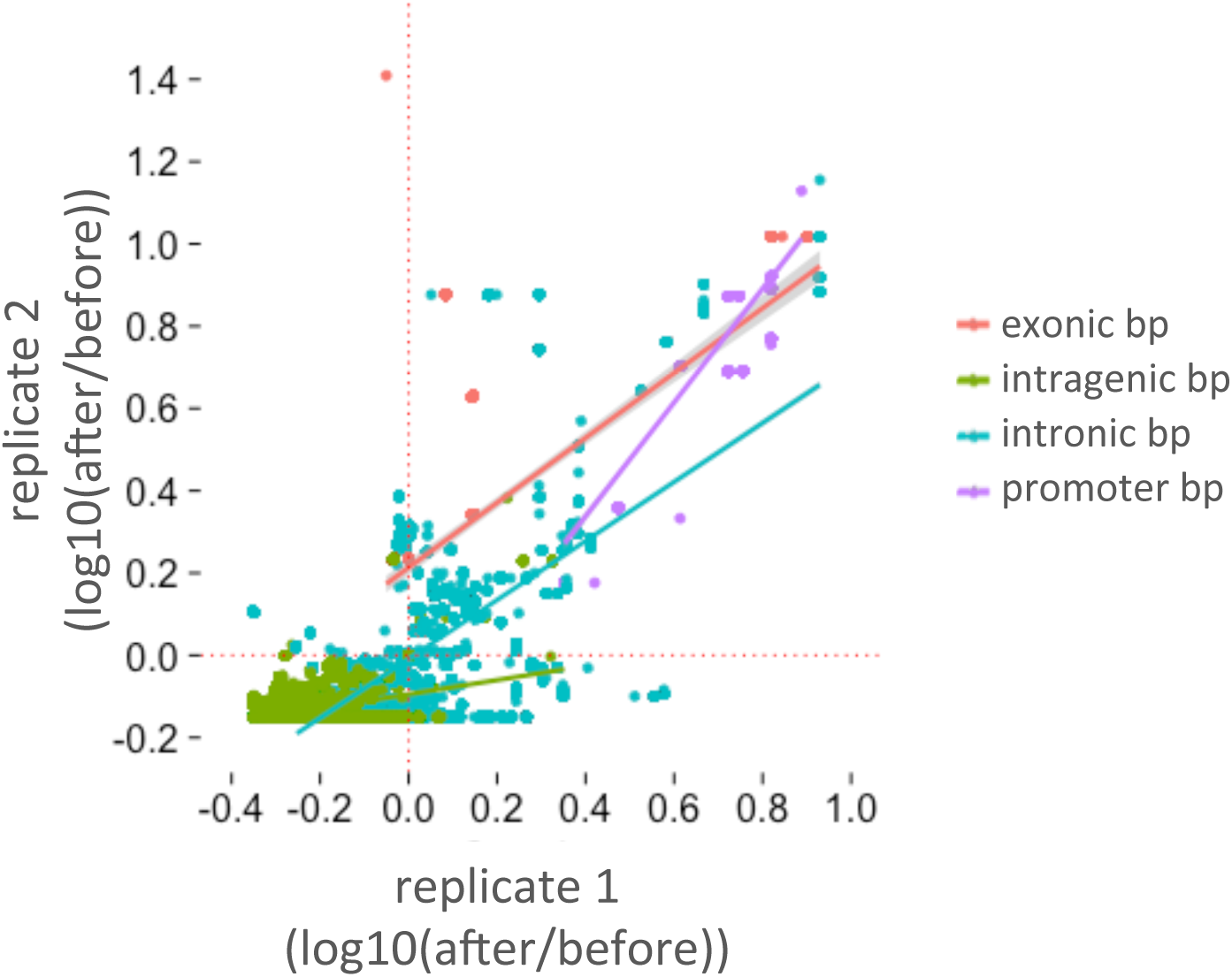
ScanDel scores correlate across two biological replicates. The ScanDel selection scores for each biological replicate were calculated per base-pair by averaging the log10(after/before 6TG) for every programmed deletion that covers that base-pair. Least squares lines and points are colored by sequence content category. The stronger correlation for the ‘intronic’ category is driven by sequences proximal to the exons as seen in Fig. 3. Red corresponds to exons (Pearson: 0.736); green to intragenic regions (Pearson: 0.417); blue to intronic regions (within 2 Kb of an exon, Pearson: 0.628; deeply intronic, Pearson: −0.0194); and purple is the promoter (1 Kb upstream of the TSS, Pearson: 0.905).

**Supplementary Figure 4:**
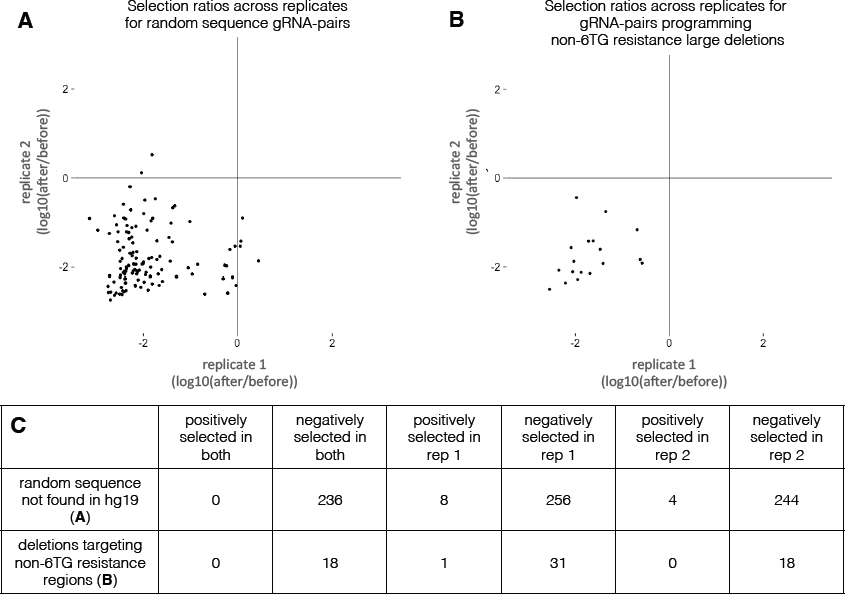
None of the negative control gRNA pairs were positively selected by 6TG in both ScanDel replicates. **A)** Negative control gRNA pairs targeting random sequences not found in hg19 were given a selection score of log10(after/before 6TG). Only gRNA pairs sampled in both replicates are plotted. **B)** Additional negative control gRNA pairs were programmed to create 1 and 2 Kb deletions in regions not expected to cause 6TG resistance. Selection scores were calculated for each gRNA pair as in **A**, and plotted for gRNA pairs found in both replicates. These region’s coordinates were randomly generated from poorly conserved sequence (Pollard, Hubisz, Rosenbloom, & Siepel, 2010) not within 10 Kb of any gene and far from *HPRT1* (chr8:23768553-23771053, chr4:25697737-25700237, chr9:41022164-41024664, chr5:12539119-12541619, chr6:23837183-23839683, chr8:11072736-11075236). **C)** Table showing counts of positively and negatively selected negative control gRNA pairs across experiments.

**Supplementary Figure 5:**
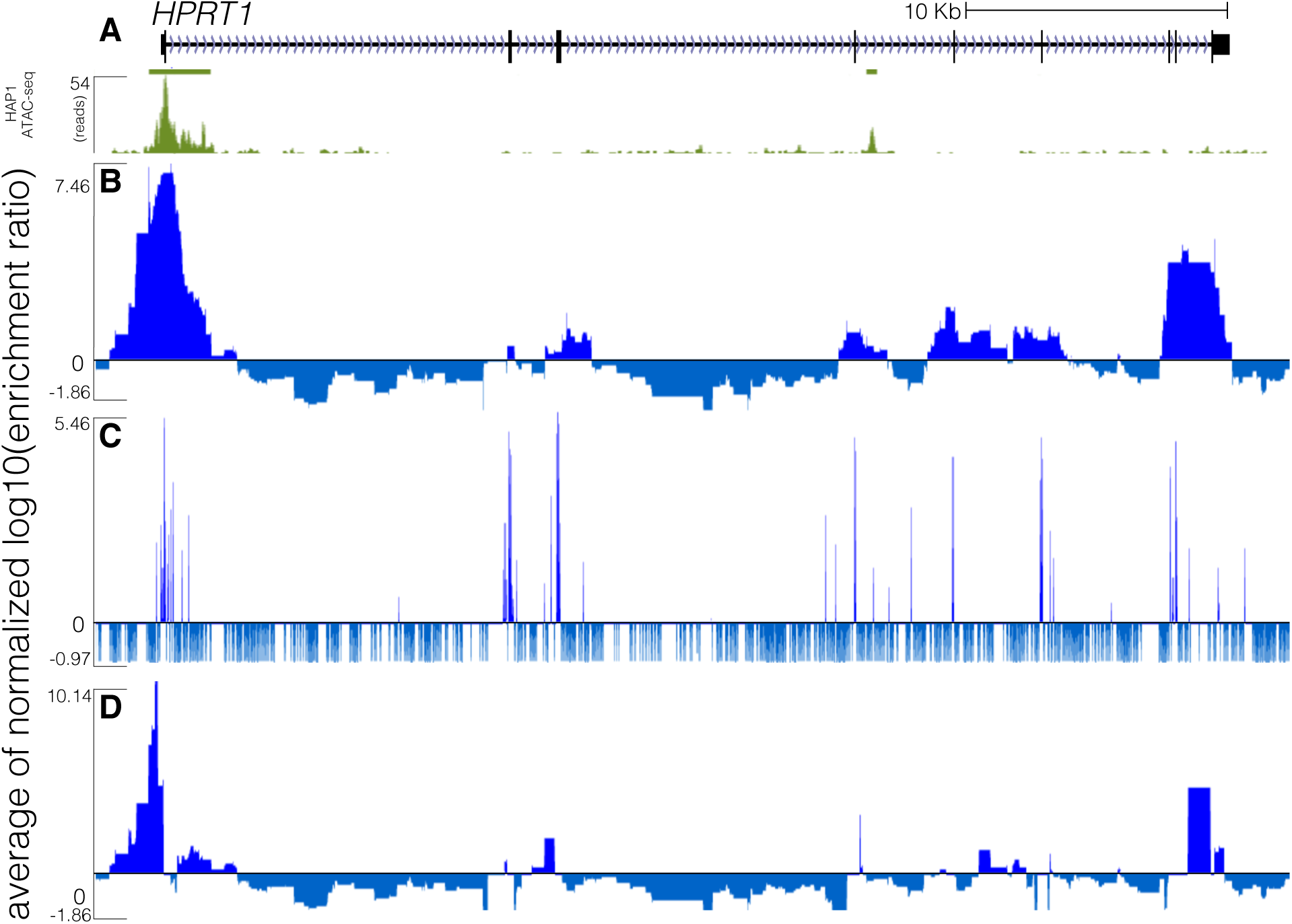
All exons and some exon-proximal non-coding regions score strongly in both the ScanDel gRNA pair screen and the individual gRNA screen. **A)** ATAC-seq data (green) from the HAP1 cell line displayed for the *HPRT1* locus ((chrX:133,591,675-133,637,198, hg19). Bars depict hotspots (John et al., 2011) and beneath is the pile-up representation of ATAC-seq reads. **B)** The same ScanDel data is displayed as in Fig. 2C but zoomed-in on the *HPRT1* locus. Each base-pair’s score is the mean of the log10(after/before 6TG) values for all the programmed deletions that cover that base-pair. These scores are normalized to the median positive score from the replicate. The average of the two replicates’ scores for each base-pair is displayed. **C)** The same individual gRNA data is displayed as in Fig. 2D but zoomed in on *HPRT1*. Each base-pair score is the mean of the log10(after/before 6TG) values for all the inferred ∼10 bp deletions that remove that base-pair. The normalized average of the two replicates’ scores for that base-pair is displayed. **D)** The same ScanDel track as in **A** but with per base-pair scores calculated after excluding any deletions programmed to disrupt an exon.

**Supplementary Figure 6:**
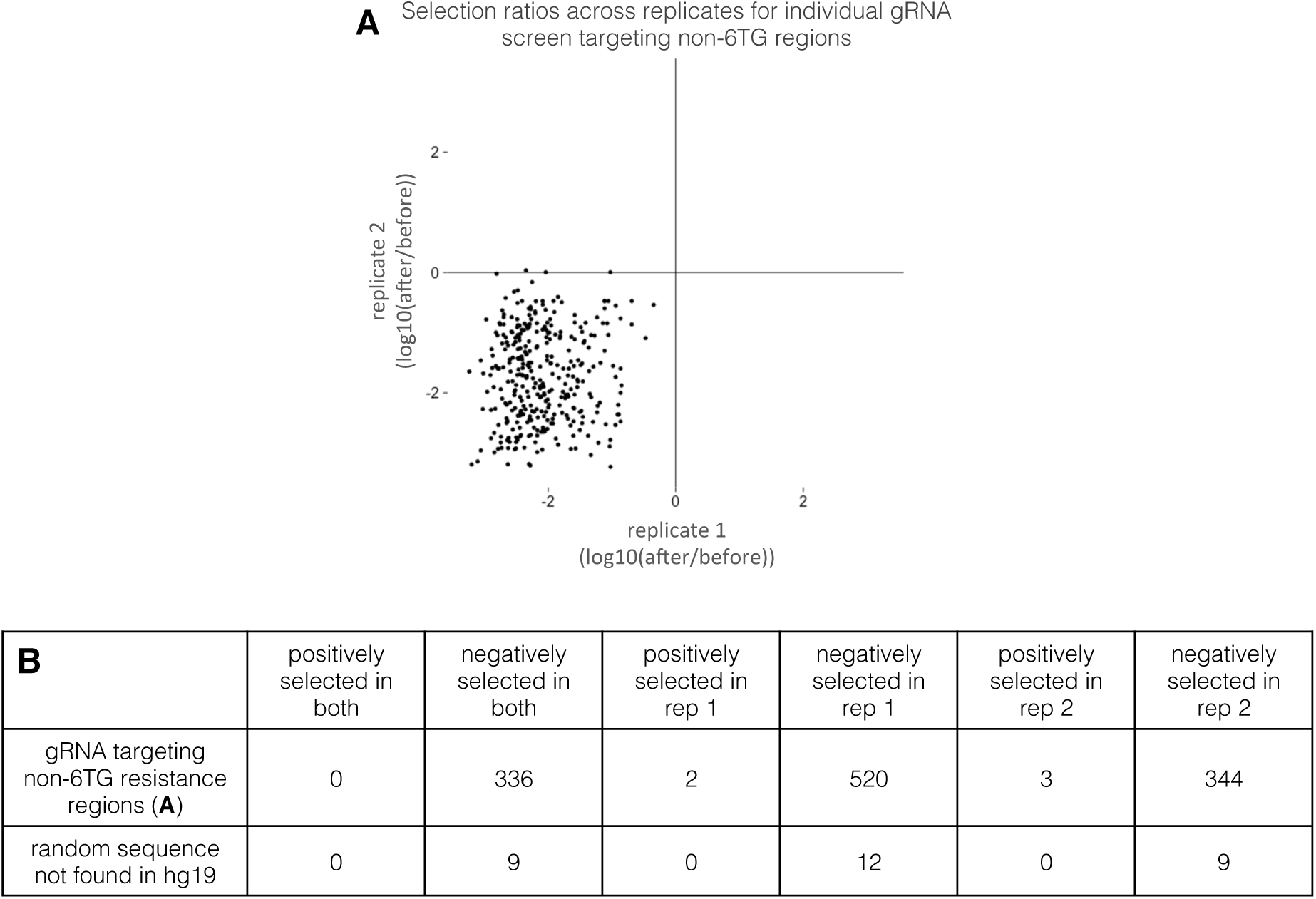
None of the negative control random-sequence gRNAs were positively selected in both individual gRNA screen replicates. **A)** Selection scores across replicates for individual gRNAs that target regions not expected to induce 6TG resistance (as in Supplementary Figure 4). Only gRNAs sampled in both replicates are plotted. **B)** Table of the negative control gRNAs selected in both, either, or neither biological replicate.

**Supplementary Figure 7:**
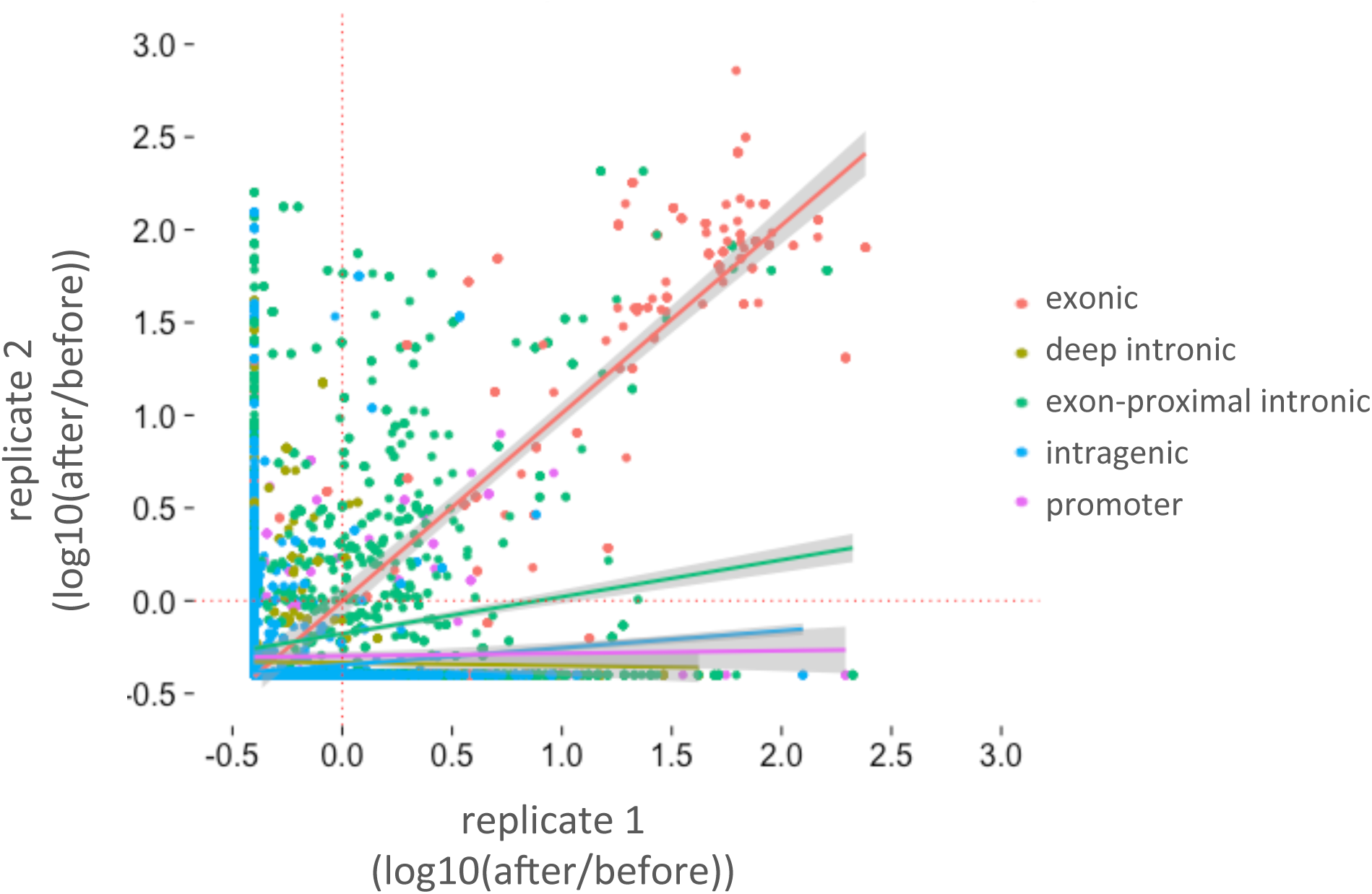
Correlation of the individual gRNA screen scores across two biological replicates. The individual gRNA scores for each biological replicate were calculated per base-pair and presented as mean of log10(after/before 6TG) between replicates. Least squares lines and points are colored by sequence content category. Specifically, intronic sequence within 2 Kb of an exon is colored in green (Pearson: 0.176); exons are red (Pearson: 0.818); deep intronic is yellow (Pearson: −0.14); intragenic sequences are blue (Pearson: 0.070; and promoter sequence (2 Kb upstream of the TSS) is purple (Pearson: 0.022).

**Supplementary Figure 8:**
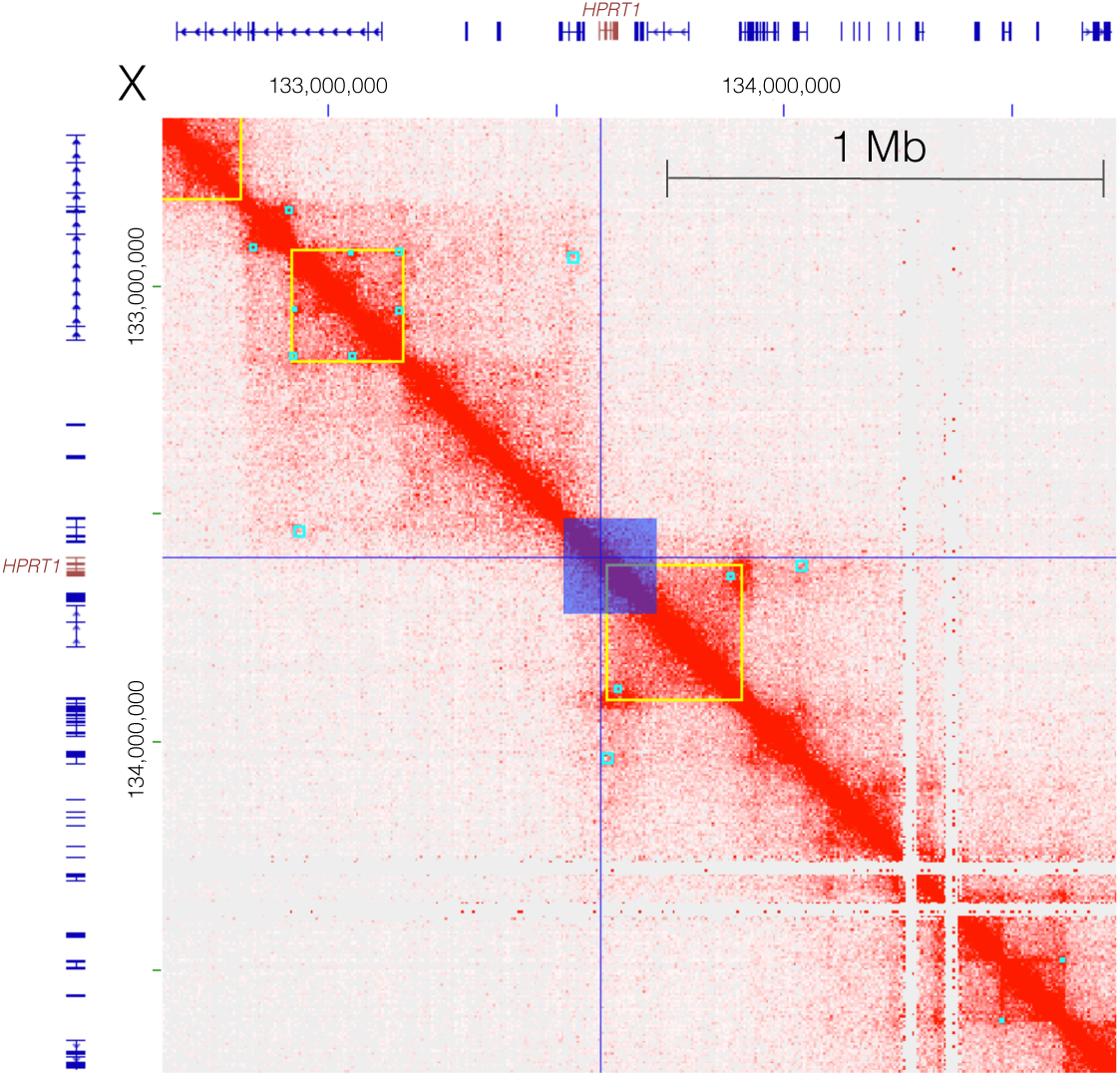
Region interrogated with ScanDel only partially surveys a 300 Kb topologically associated domain (TAD) found in HAP1 cells. A heatmap of interactions between 5 Kb bins along chrX:132,669,000-134,716,000 (hg19) in HAP1 cells (Sanborn et al., 2015) (Juicebox 1.4 (Durand et al., 2016), balanced normalization). RefSeq gene annotations are drawn across the axes, with the *HPRT1* gene model drawn in red. Blue lines mark its TSS and the 206 Kb surveyed by ScanDel is highlighted as a dark blue box. Light blue boxes mark peaks and yellow boxes mark TADs as called by Sanborn et al.

**Supplementary Figure 9:**
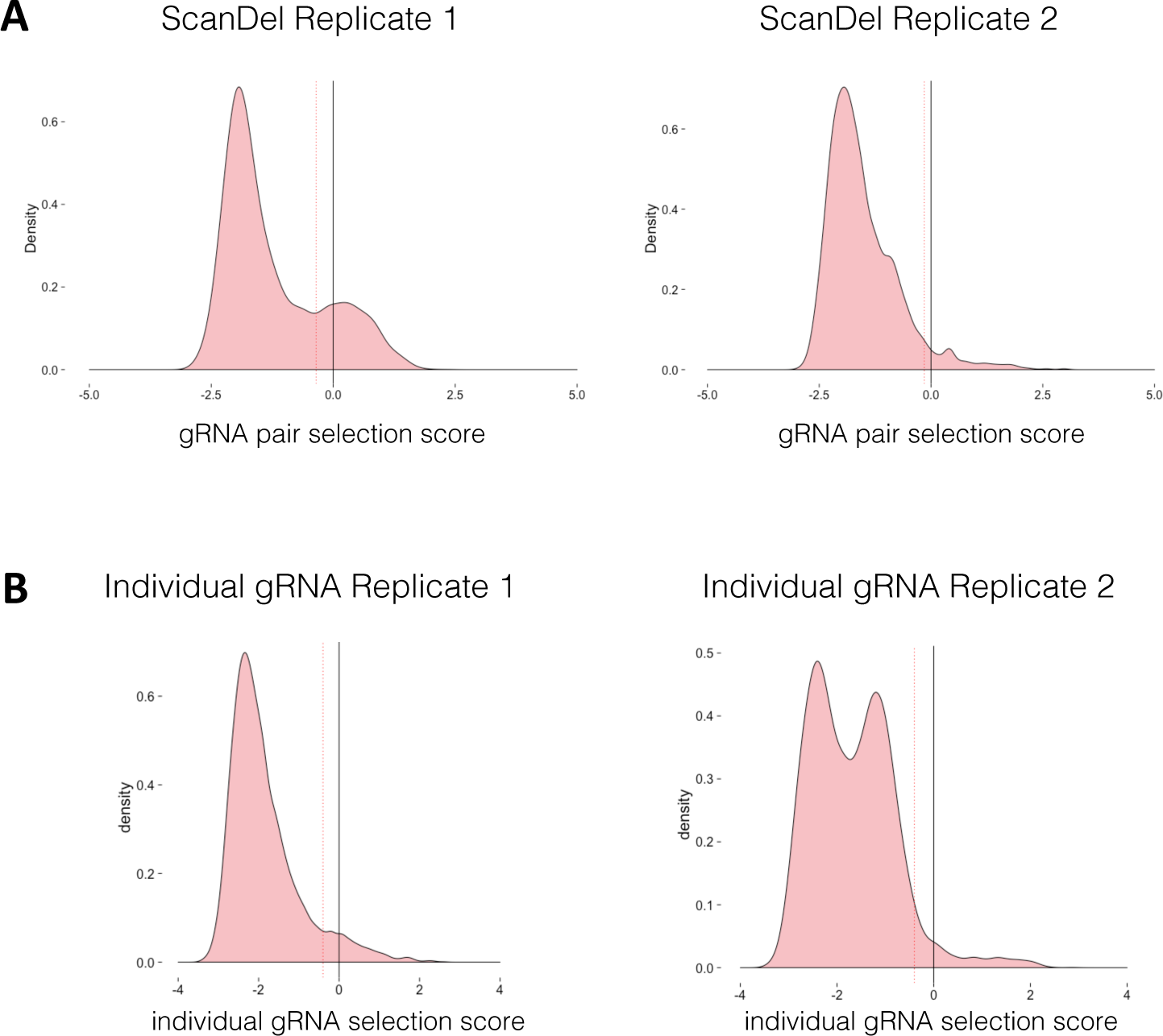
Distribution of selection scores across biological replicates for ScanDel gRNA pairs or individual gRNAs. **A)** Each gRNA pair in the ScanDel screens was assigned a selection score (log10(after/before 6TG)). The minimum selection score threshold described in Methods (-0.35 for replicate 1, −0.15 for replicate 2) is drawn with a dotted red line. **B)** Each gRNA in the individual gRNA screen was assigned a selection score as in A, for each replicate. The minimum negative selection score threshold (−0.4 for both replicates) is drawn with a dotted red line (explanation in Methods).

**Supplementary Table 1.**
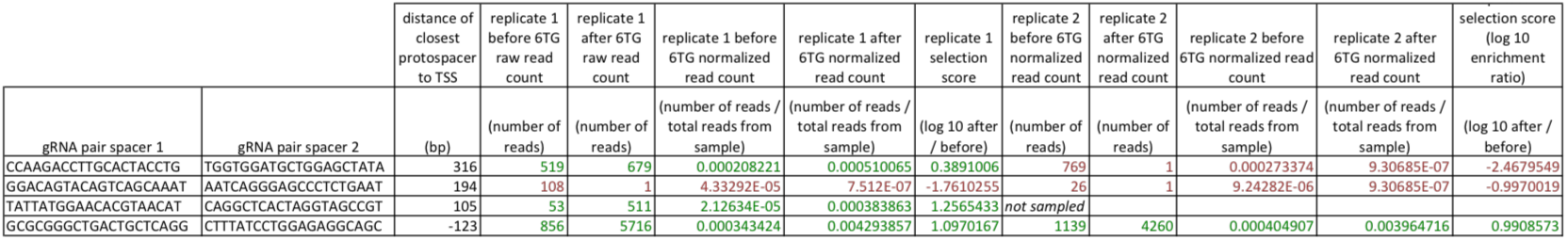
Read count data and selection scores for the 4 gRNA pairs upstream of exon 1 used for Fig. 3A-D. Green is positively selected and red is negatively selected

**Supplementary Table 2.**
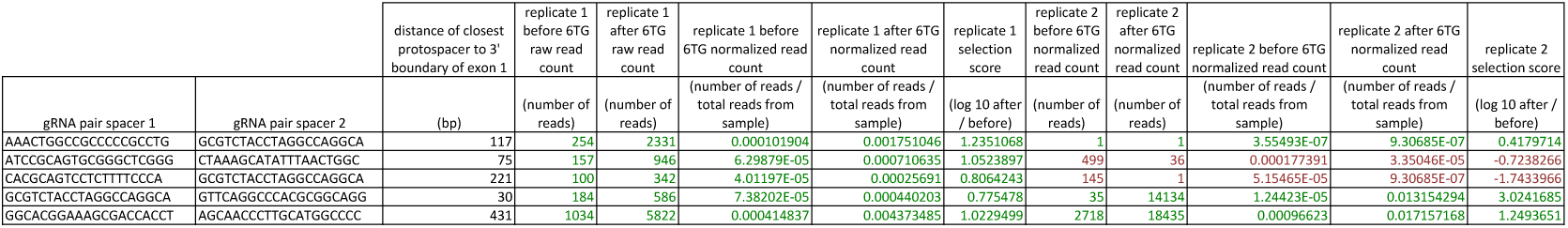
Read count data and selection scores for the 5 gRNA pairs in intron 1 used for Fig. 3E-H

**Supplementary Table 3.**
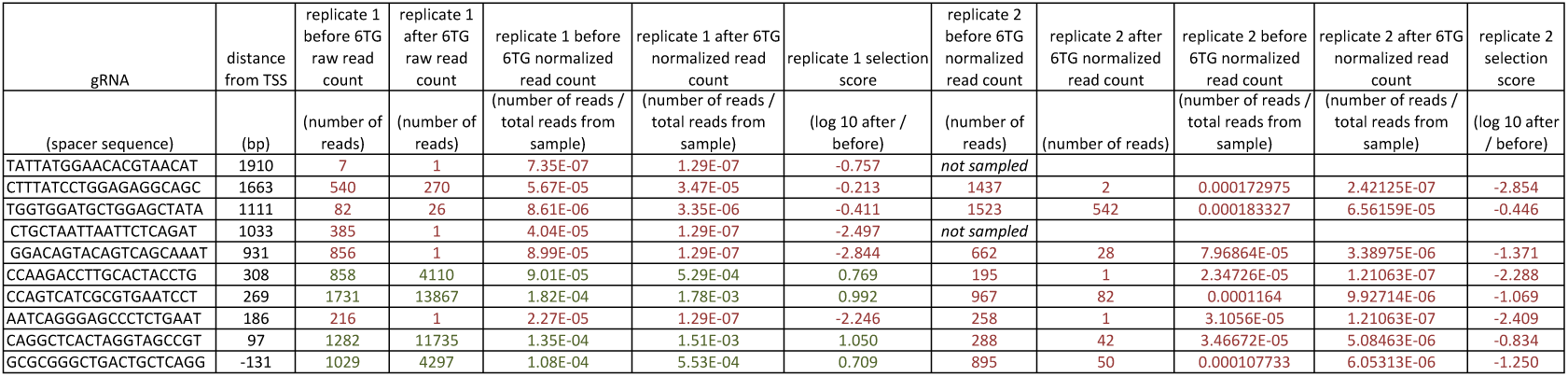
Read count data and selection scores (for both replicates of the individual gRNA screen) for the 10 individual gRNAs targeting regions upstream of exon 1 and displayed in Fig. 4.

## References

Aguet, F., Brown, A. A., Castel, S., Davis, J. R., Mohammadi, P., Segre, A. V, … Montgomery, S. B. (2016). Local genetic effects on gene expression across 44 human tissues. bioRxiv.

Aparicio-Prat, E., Arnan, C., Sala, I., Bosch, N., Guigó, R., & Johnson, R. (2015). DECKO: Single-oligo, dual-CRISPR deletion of genomic elements including long non-coding RNAs. BMC Genomics, 16(1), 846. http://doi.org/10.1186/s12864-015-2086-z

Ardlie, K. G., Deluca, D. S., Segre, A. V., Sullivan, T. J., Young, T. R., Gelfand, E. T., … Lockhart, N.C. (2015). The Genotype-Tissue Expression (GTEx) pilot analysis: Multitissue gene regulation in humans. Science, 348(6235), 648–660. http://doi.org/10.1126/science.1262110

Banerji, J., Rusconi, S., & Schaffner, W. (1981). Expression of a beta-globin gene is enhanced by remote SV40 DNA sequences. Cell, 27(2 PART 1), 299–308. http://doi.org/10.1016/0092-8674(81)90413-X

Bolger, A. M., Lohse, M., & Usadel, B. (2014). A flexible trimmer for Illumina sequence data. Bioinformatics, 30, 2114.

Boyle, A. P., Hong, E. L., Hariharan, M., Cheng, Y., Schaub, M. A., Kasowski, M., … Snyder, M. (2012). Annotation of functional variation in personal genomes using RegulomeDB. Genome Research, 22(9), 1790–1797. http://doi.org/10.1101/gr.137323.112

Buenrostro, J. D., Giresi, P. G., Zaba, L. C., Chang, H. Y., & Greenleaf, W. J. (2013). Transposition of native chromatin for fast and sensitive epigenomic profiling of open chromatin, DNA-binding proteins and nucleosome position. Nat Meth, 10(12), 1213–1218. JOUR. Retrieved from http://dx.doi.org/10.1038/nmeth.2688

Byrne, S. M., Ortiz, L., Mali, P., Aach, J., & Church, G. M. (2015). Multi-kilobase homozygous targeted gene replacement in human induced pluripotent stem cells. Nucleic Acids Research, 43(3), e21. http://doi.org/10.1093/nar/gku1246

Canver, M. C., Bauer, D. E., Dass, A., Yien, Y. Y., Chung, J., Masuda, T., … Orkin, S. H. (2014). Characterization of genomic deletion efficiency mediated by clustered regularly interspaced palindromic repeats (CRISPR)/cas9 nuclease system in mammalian cells. Journal of Biological Chemistry, 289(31), 21312–21324. http://doi.org/10.1074/jbc.M114.564625

Canver, M. C., Smith, E. C., Sher, F., Pinello, L., Sanjana, N. E., Shalem, O., … Bauer, D. E. (2015). BCL11A enhancer dissection by Cas9-mediated in situ saturating mutagenesis. Nature. http://doi.org/10.1038/nature15521

Chen, S., Sanjana, N. E., Zheng, K., Shalem, O., Lee, K., Shi, X., … Sharp, P. A. (2015). Genome-wide CRISPR screen in a mouse model of tumor growth and metastasis. Cell, 160(6), 1246–1260. http://doi.org/10.1016/j.cell.2015.02.038

Chong, J. X., Buckingham, K. J., Jhangiani, S. N., Boehm, C., Sobreira, N., Smith, J. D., … Bamshad, M. J. (2015). The Genetic Basis of Mendelian Phenotypes: Discoveries, Challenges, and Opportunities. American Journal of Human Genetics, 97(2), 199–215. http://doi.org/10.1016/j.ajhg.2015.06.009

Coetzee, S. G., Rhie, S. K., Berman, B. P., Coetzee, G. A., & Noushmehr, H. (2012). FunciSNP: An R/bioconductor tool integrating functional non-coding data sets with genetic association studies to identify candidate regulatory SNPs. Nucleic Acids Research, 40(18). http://doi.org/10.1093/nar/gks542

Diao, Y., Li, B., Meng, Z., Jung, I., Lee, A., Dixon, J., … Ren, B. (2016). A new class of temporarily phenotypic enhancers identified by CRISPR/Cas9 mediated genetic screening. Genome Research, 1–9. http://doi.org/10.1101/gr.197152.115

Doench, J. G., Fusi, N., Sullender, M., Hegde, M., Vaimberg, E. W., Donovan, K. F., … Root, D. E. (2016). Optimized sgRNA design to maximize activity and minimize off-target effects of CRISPR-Cas9. Nature Biotechnology, 34(November 2015), 1–12. http://doi.org/10.1038/nbt.3437

Döring, A., Weese, D., Rausch, T., & Reinert, K. (2008). SeqAn an efficient, generic C++ library for sequence analysis. BMC Bioinformatics, 9, 11. http://doi.org/10.1186/1471-2105-9-11

Durand, N. C., Robinson, J. T., Shamim, M. S., Machol, I., Mesirov, J. P., Lander, E. S., & Aiden, E. L. (2016). Juicebox Provides a Visualization System for Hi-C Contact Maps with Unlimited Zoom. Cell Systems, 3(1), 99–101. http://doi.org/10.1016Zj.cels.2015.07.012

ENCODE Project Consortium, Bernstein, B. E., Birney, E., Dunham, I., Green, E. D., Gunter, C., & Snyder, M. (2012). An integrated encyclopedia of DNA elements in the human genome. Nature, 489(7414), 57–74. http://doi.org/nature11247[pii]/n10.1038/nature11247

Ernst, J., & Kellis, M. (2012). ChromHMM: automating chromatin-state discovery and characterization. Nature Methods, 9(3), 215–6. http://doi.org/10.1038/nmeth.1906

Fu, R., Ceballos-Picot, L., Rosa, J., Larovere, L. E., Yamada, Y., Nguyen, K. V., … Jinnah, H. A. (2014). Genotype-phenotype correlations in neurogenetics: Lesch-Nyhan disease as a model disorder. Brain, 137(5), 1282–1303. http://doi.org/10.1093/brain/awt202

Hoffman, M. M., Buske, O. J., Wang, J., Weng, Z., Bilmes, J. a, & Noble, W. S. (2012). Unsupervised pattern discovery in human chromatin structure through genomic segmentation. Nature Methods, 9(5), 473–6. http://doi.org/10.1038/nmeth.1937

Hsu, P. D., Scott, D. a, Weinstein, J. a, Ran, F. A., Konermann, S., Agarwala, V., … Zhang, F. (2013). DNA targeting specificity of RNA-guided Cas9 nucleases. Nature Biotechnology, 31(9), 827–32. http://doi.org/10.1038/nbt.2647

John, S., Sabo, P. J., Thurman, R. E., Sung, M.-H., Biddie, S. C., Johnson, T. A., … Stamatoyannopoulos, J. A. (2011). Chromatin accessibility pre-determines glucocorticoid receptor binding patterns. Nature Genetics, 43(3), 264–268. http://doi.org/10.1038/ng.759

Kircher, M., Witten, D. M., Jain, P., O’Roak, B. J., Cooper, G. M., & Shendure, J. (2014). A general framework for estimating the relative pathogenicity of human genetic variants. Nature Genetics, 46(3), 310–315. http://doi.org/10.1038/ng.2892

Korkmaz, G., Lopes, R., Ugalde, A. P., Nevedomskaya, E., Han, R., Myacheva, K., … Agami, R. (2016). Functional genetic screens for enhancer elements in the human genome using CRISPR-Cas9. Nature Biotechnology, (August 2015), 1–10. http://doi.org/10.1038/nbt.3450

Langmead, B., & Salzberg, S. L. (2012). Fast gapped-read alignment with Bowtie 2. Nat Methods, 9(4), 357–359. http://doi.org/10.1038/nmeth.1923

Lesch, M., & Nyhan, W. L. W. (1964). A familial disorder of uric acid metabolism and central nervous system function. The American Journal of Medicine, 9(April), 561–570. http://doi.org/10.1016/0002-9343(64)90104-4

Li, M. J., Wang, L. Y., Xia, Z., Sham, P. C., & Wang, J. (2013). GWAS3D: Detecting human regulatory variants by integrative analysis of genome-wide associations, chromosome interactions and histone modifications. Nucleic Acids Research, 41(Web Server issue), 150–158. http://doi.org/10.1093/nar/gkt456

Maurano, M. T., Humbert, R., Rynes, E., Thurman, R. E., Haugen, E., Wang, H., … Stamatoyannopoulos, J. A. (2012). Systematic Localization of Common Disease-Associated Variation in Regulatory DNA. Science, 337(6099), 1190–1195. http://doi.org/10.1126/science.1222794

McKenna, A., Findlay, G. M., Gagnon, J. A., Horwitz, M. S., Schier, A. F., & Shendure, J. (2016). Whole-organism lineage tracing by combinatorial and cumulative genome editing. Science, 353(6298), aaf7907–aaf7907. http://doi.org/10.1126/science.aaf7907

Patwardhan, R. P., Lee, C., Litvin, O., Young, D. L., Pe’er, D., & Shendure, J. (2009). High-resolution analysis of DNA regulatory elements by synthetic saturation mutagenesis. Nature Biotechnology, 27(12), 1173–1175. http://doi.org/10.1038/nbt.1589

Pollard, K. S., Hubisz, M. J., Rosenbloom, K. R., & Siepel, A. (2010). Detection of nonneutral substitution rates on mammalian phylogenies. Genome Research, 20(1), 110–121. http://doi.org/10.1101/gr.097857.109

Rajagopal, N., Srinivasan, S., Kooshesh, K., Guo, Y., Edwards, M. D., Banerjee, B., … Sherwood, R. I. (2016). High-throughput mapping of regulatory DNA. Nature Biotechnology, (January). http://doi.org/10.1038/nbt.3468

Reid, L. H., Gregg, R. G., Smithies, O., & Koller, B. H. (1990). Regulatory elements in the introns of the human HPRT gene are necessary for its expression in embryonic stem cells. Proceedings of the National Academy of Sciences of the United States of America, 87(11), 4299–303. http://doi.org/10.1073/pnas.87.11.4299

Rice, P., Longden, I., & Bleasby, A. (2000). EMBOSS: The European Molecular Biology Open Software Suite. Trends in Genetics, 16(1), 276–277. http://doi.org/10.1016/j.cocis.2008.07.002

Rincon-Limas, D. E., & Krueger, D. A. (1991). Functional Characterization of the Human Hypoxanthine Phosphoribosyltransferase Gene Promoter: Evidence for a Negative Regulatory Element. Molecular and Cellular Biology, 11(8), 4157–4164.

Sanborn, A. L., Rao, S. S. P., Huang, S.C., Durand, N. C., Huntley, M. H., Jewett, A. I., … Aiden, E. L. (2015). Chromatin extrusion explains key features of loop and domain formation in wild-type and engineered genomes. Proceedings of the National Academy of Sciences, 112(47), 201518552. http://doi.org/10.1073/pnas.1518552112

Sanjana, N. E., Shalem, O., & Zhang, F. (2014). Improved vectors and genome-wide libraries for CRISPR screening. Nature Methods, 11(8), 783–784. http://doi.org/10.1038/nmeth.3047

Sanjana, N. E., Wright, J., Zheng, K., Shalem, O., Fontanillas, P., Joung, J., … Zhang, F. (2016). High-resolution interrogation of functional elements in the noncoding genome. Science, 353(6307), 1545–1549.

Shalem, O., Sanjana, N. E., Hartenian, E., Shi, X., Scott, D. A., Mikkelsen, T. S., … Zhang, F. (2014). Genome-scale CRISPR-Cas9 knockout screening in human cells. Science, 343(6166), 1–78. http://doi.org/10.1126/science.1247005.Genome-Scale

Tsai, S. Q., Zheng, Z., Nguyen, N. T., Liebers, M., Topkar, V. V, Thapar, V., … Joung, J. K. (2014). GUIDE-seq enables genome-wide profiling of off-target cleavage by CRISPR-Cas nucleases. Nature Biotechnology, 33(2), 187–198. http://doi.org/10.1038/nbt.3117

Wang, T., Wei, J. J., Sabatini, D. M., & Lander, E. S. (2014). Genetic screens in human cells using the CRISPR-Cas9 system. Science (New York, N.Y.), 343(6166), 80–4. http://doi.org/10.1126/science.1246981

Ward, L. D., & Kellis, M. (2012). HaploReg: A resource for exploring chromatin states, conservation, and regulatory motif alterations within sets of genetically linked variants. Nucleic Acids Research, 40(D1), 930–934. http://doi.org/10.1093/nar/gkr917

Zhou, Y., Zhu, S., Cai, C., Yuan, P., Li, C., Huang, Y., & Wei, W. (2014). High-throughput screening of a CRISPR/Cas9 library for functional genomics in human cells. Nature, 509(7501), 487–91. http://doi.org/10.1038/nature13166

Zhu, S., Li, W., Liu, J., Chen, C.H., Liao, Q., Xu, P., … Wei, W. (2016). Genome-scale deletion screening of human long non-coding RNAs using a paired-guide RNA CRISPR–Cas9 library. Nature Biotechnology, 2016(October). http://doi.org/10.1038/nbt.3715

